# PRISM: ancestry-aware integration of tissue-specific genomic annotations enhances the transferability of polygenic scores

**DOI:** 10.1101/2025.11.13.688144

**Authors:** Xiaohe Tian, Tabassum Fabiha, William F Li, Kushal K Dey, Manolis Kellis, Yosuke Tanigawa

## Abstract

The limited transferability of polygenic scores (PGS) across populations constrains their clinical utility and risks exacerbating health disparities, given challenges in multi-ancestry training, fine-mapping, and variant prioritization using genomic annotations, particularly when biologically relevant reference resources are sparse or unavailable for the target population. Here, we introduce PRISM, a transfer learning approach that jointly addresses these challenges to enhance PGS transferability. Applying PRISM to 7352 fine-mapped variants, 414 ENCODE annotations, and 406,659 individuals from the UK Biobank, we demonstrate that ancestry-aware integration of tissue-specific annotations yields the largest gains in predictive performance for African ancestry, with an average improvement of 13.10% (p=1.6×10^−5^) over annotation-agnostic multi-ancestry PGS. Notably, the best-performing model uses 102-fold fewer annotations than non-specific models, with contributions from broad categories of annotations. Overall, PRISM complements ongoing data diversification efforts by providing an immediately applicable strategy based on the integration of biologically aligned, best-available resources to address genomic health equity.

## Introduction

Translating genomic discoveries into accurate predictive models of genetic liability is one of the central goals of human genetics, with substantial implications for disease risk prediction, risk stratification, and population health research. Advances in genome-wide association studies (GWAS) have identified thousands of loci associated with complex traits and diseases, fueling methodological developments and their applications for estimating polygenic contributions to disease liability. Polygenic scores (PGS), a statistical approach that aggregates genome-wide genetic effects into a single numeric value, have attracted substantial research interest and are increasingly explored for clinical applications[1,2]. However, the limited transferability of PGS across populations and its potential risks for exacerbating health disparities remain one of the most significant challenges[3,4]: PGS models trained in one training cohort often underperform in other target populations, especially in African ancestry populations[5], which harbor substantial genetic diversity but often exhibit the lowest predictive accuracy[6–9]. In addition to the ongoing data collection and capacity-building efforts, such as those led by H3Africa[10–13], there is a pressing need for computational and statistical approaches that enhance PGS performance in underrepresented populations by maximizing the value of currently available resources[14].

Three main computational strategies have emerged to address the limited transferability of polygenic scores. First, integration of genomic annotations, such as atlases of putative regulatory elements or cell-type-specific transcription factor binding patterns, helps prioritize variants with functional relevance[15,16]. Second, training PGS with multi-ancestry data leverages the shared genetic architecture of complex traits across genetic ancestry groups, effectively prioritizing more robust genetic effects through pooling and implicit replication across populations[17,18]. Recent work extends this strategy by training across the continuum of genetic ancestry, as in our previous inclusive PGS (iPGS)[18], which demonstrated improvements across all ancestry groups. Third, statistical fine-mapping prioritizes likely causal variants by resolving correlated association signals in regions of high linkage disequilibrium (LD)[19]. Some methods incorporate combinations of these strategies[20,21], but how to effectively integrate large-scale genomic annotations, multi-ancestry modeling, and statistical fine-mapping that maximize benefits given limited and uneven data availability across populations and tissues is still largely unexplored.

The increasing availability of genomic annotations presents a particularly attractive opportunity for enhancing polygenic score modeling by prioritizing likely causal variants. For example, the ENCODE Consortium has generated one of the largest and most comprehensive collections of regulatory annotations to date, consisting of both experimentally derived and computationally predicted genomic annotation tracks across a wide range of tissues and cell types[22–25]. These resources enable the integration of regulatory genomics into statistical genetics, as demonstrated in pioneering works[15,20,21]. However, several practical challenges remain underexplored: it is unclear which annotation modalities are most informative for a given trait or population, how best to combine potentially heterogeneous annotations, and how to proceed when trait-relevant annotations are not readily available for the target population of interest[26]. Developing new methodologies to address these open questions would enhance the transferability of PGS, especially for underrepresented populations where statistical power is often limited.

Here, we overcome these technical limitations and present Priors-informed Regression for Inclusive Score Modeling (PRISM), a transfer learning based framework that unifies three complementary strategies for enhancing PGS transferability: variant prioritization using large-scale genomic annotations, multi-ancestry modeling, and statistical fine-mapping. With PRISM, we apply transfer learning to derive variant-level scores from large-scale annotations and fine-mapped variants, which we subsequently use to inform variant prioritization in polygenic modeling. We apply PRISM to integrate 414 genomic annotations from the ENCODE Phase IV release with 7352 ancestry-specific, trait-agnostic fine-mapped variants from the Million Veteran Program (MVP) and use these scores to train PGS models using 406,659 individuals across the continuum of genetic ancestry in the UK Biobank (UKB)[11,27,28]. We demonstrate that ancestry-aware integration of tissue-specific annotations leads to the greatest improvements in PGS transferability, with an average improvement of 13.1% (p=1.6×10^−5^) in *R*^2^ in UKB African ancestry individuals across select traits, despite the limited availability of 102-fold fewer biologically aligned genomic annotations and only 1.49% of ancestry-matched individuals used in the PGS training. PRISM also facilitates biological interpretation by revealing heterogeneous contributions of trait-relevant annotations. Overall, our work highlights the advantage of integrating biologically aligned annotations as a complementary strategy to enhance PGS transferability, offering pragmatic and immediately applicable approaches to advance precision health for all.

## Results

### Overview of the PRISM and study design

In PRISM, we integrate large-scale annotations with fine-mapping results through transfer learning to enhance PGS transferability (**Fig. 1a**). The workflow consists of three main steps, whose modular design allows us to consider ancestry at multiple components of the analysis. First, we curate annotations and fine-mapping results across a large number of traits. Second, we train a supervised learning model using fine-mapping results as labels and annotations as input features, resulting in variant-level annotation scores. These scores capture the optimal combination of annotations supporting the biological relevance of variants. Finally, we use the scores to prioritize variants in PGS training.

**Figure 1.**
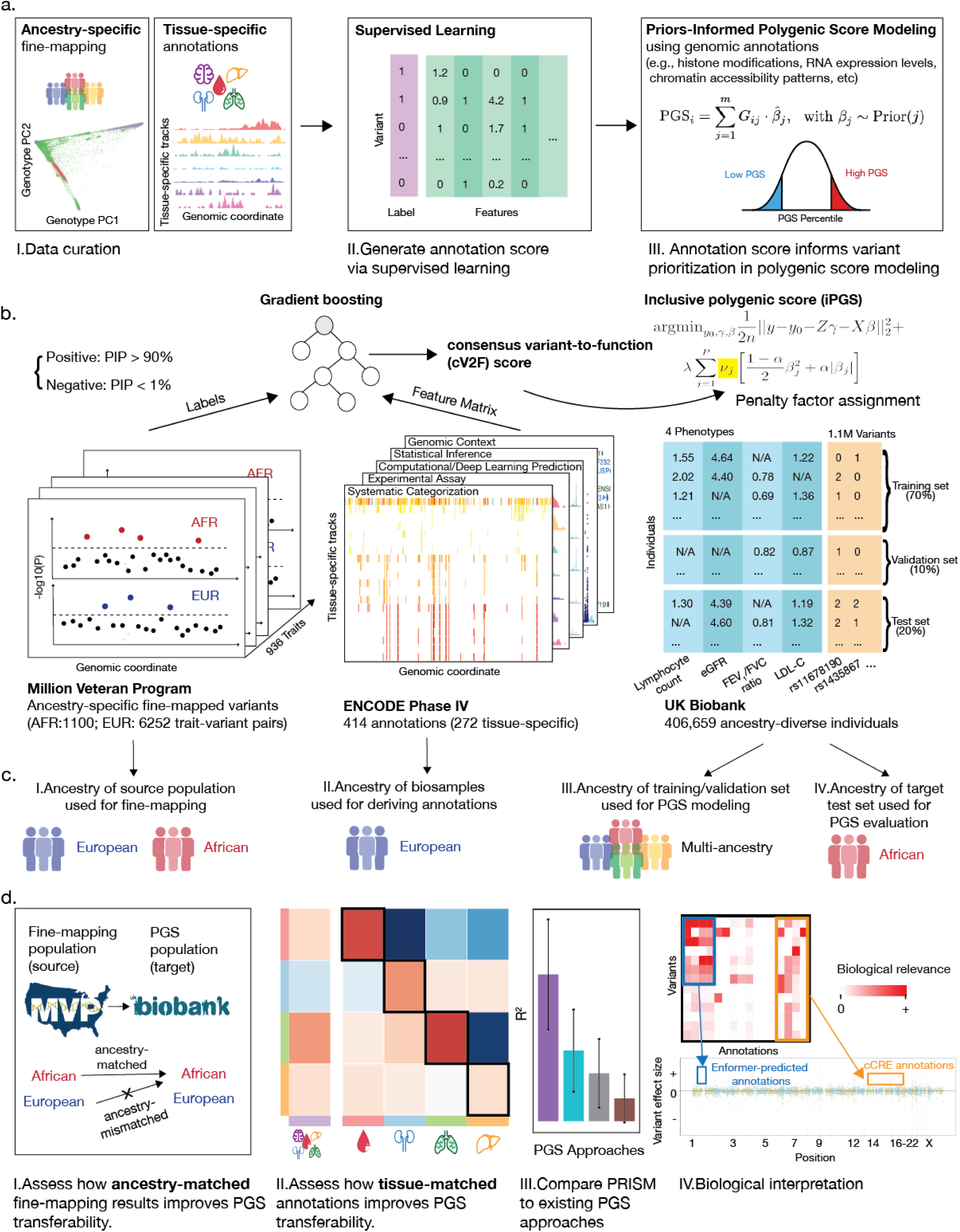
Overview of the PRISM and study design. **a.** PRISM (Priors-informed Regression for Inclusive Score Modeling using genomic annotations) integrates genomic annotations with fine-mapping results to inform polygenic score (PGS) development via transfer learning. The workflow consists of three steps: (I) curation of annotations and fine-mapping results, (II) supervised learning to generate annotation-informed scores, and (III) PGS training with variant prioritization. **b.** Application of PRISM in this study. We integrated annotations from ENCODE, ancestry-specific fine-mapping results from the Million Veteran Program (MVP), and genetic and phenotypic data from the UKB to train PGS models across four traits with known primary tissues. We aggregated annotations into consensus variant-to-function (cV2F) scores using gradient boosting and utilized them in inclusive PGS (iPGS). **c.** PRISM accounts for four types of ancestry in PGS development: (I) the fine-mapping source population, (II) the ancestry of biosamples used to generate annotations, (III) the ancestry of individuals in PGS model development, and (IV) the ancestry of the held-out test set used for evaluation. **d.** Systematic comparisons and interpretation. We evaluated the effects of ancestry-matched fine-mapping and tissue-matched annotations on PGS transferability. We compared PRISM to existing PGS approaches. We performed biological interpretation to examine the annotations contributing to improved predictive performance in PRISM.

In our study, we applied PRISM to annotations from ENCODE, fine-mapping results from MVP, and individual-level genetic and phenotypic data from the UKB (**Fig. 1b**). Specifically, we first curated ancestry-specific fine-mapped variants from MVP (7352 trait-variant pairs across 936 traits) and 414 annotations from ENCODE (**Data and code availability**)[11,22,29–33]. Second, we applied gradient boosting to train a predictive model of continuous cV2F scores from annotations and fine-mapping results (**Methods**)[34]. Third, we used these scores in PGS training in UKB[18,28,34,35], focusing on the following four traits with clear primary tissue (**Methods**, **Supplementary Table 1**): lymphocyte count (blood), estimated glomerular filtration rate (eGFR) (kidney), forced expiratory volume in one second to forced vital capacity (FEV_1_/FVC ratio) (lung), and low-density lipoprotein cholesterol (LDL-C) (liver)[36]. We evaluated the predictive performance of PRISM models and compared them to a baseline model trained without annotations or fine-mapping results (**Methods**).

The modular design of PRISM accounts for four distinct types of ancestry in PGS modeling (**Fig. 1c**): (I) the genetic ancestry of the population used in fine-mapping (**source** population), (II) the ancestry of biosamples used to generate annotations, (III) the ancestry of individuals used for PGS development, and (IV) the ancestry of the held-out test set used for evaluation (**target** population). In this study, we focused on improving the predictive performance of the African (AFR) ancestry group in the UKB. We used either AFR or European (EUR) populations in MVP as the source for fine-mapping. For PGS training, we applied inclusive PGS (iPGS) and considered individuals across the continuum of genetic ancestry[18].

Applying PRISM, we assessed the impact of ancestry- and tissue-specific models on PGS transferability, compared its performance with existing PGS approaches, and conducted biological interpretation of the best-performing models (**Fig. 1d**). We systematically varied the source population of fine-mapping results to evaluate ancestry-matched and -mismatched PRISM models. We considered different annotation sets to evaluate tissue-matched, tissue-mismatched, and tissue-non-specific models. Tissue-matched models used annotations associated with biosamples matching the trait’s primary tissue; tissue-mismatched models used annotations from a non-primary tissue; and tissue-non-specific models used the all annotations available (**Methods**). We compared the predictive performance of PRISM to existing PGS approaches to evaluate its relative strength and limitations. To investigate how annotations contribute to variant prioritization in PRISM, we examined the annotations associated with selected variants that showed increased predictive effects.

### Ancestry-matched fine-mapping results enhance PGS transferability

To evaluate how the genetic ancestry of the source population influences PGS transferability, we trained PRISM models using fine-mapping results from individuals of either African or European ancestry in MVP and compared their performance. We quantified predictive performance (*R^2^*) in in the same held-out set of African ancestry individuals in UKB (n=1154 for lymphocyte count, n=1134 for eGFR, n=1078 for FEV_1_/FVC ratio, and n=1130 for LDL-C) and assessed average improvements over the baseline model across the four select traits using orthogonal distance regression (ODR) (**Supplementary Fig. 1**)[11,37].

We found that ancestry-matched PRISM models consistently outperformed ancestry-mismatched ones (**Table 1**). Specifically, the average improvement in predictive performance was 13.10% (p=1.6×10^−5^) for the ancestry-matched model, compared to 1.80% (p=2×10^−6^) for the ancestry-mismatched model (Model 1 vs. 2, **Methods**) when using tissue-matched annotations, despite the European cohort having 3.7 times more individuals and 5.7 times more fine-mapped variants. We also observed qualitatively similar results with tissue-non-specific annotations (Model 3 vs. 4). Overall, these results demonstrate that ancestry-matched fine-mapped variants led to more effective enhancements in PGS transferability than larger sets of ancestry-mismatched fine-mapped variants.

**Table 1.** Ancestry-matched PRISM improves PGS transferability.

|  | Annotations | Statistical fine-mapping in the source population |  |  |  |  |  | Average improvement over baseline | p-value |
| --- | --- | --- | --- | --- | --- | --- | --- | --- | --- |
| Model | Tissue specificity | Source population in fine-mapping | Ancestry-matched | Number of individuals |  | Number of the fine-mapped variants |  |  |  |
| 1 | Matched | MVP African | Yes | 121,177 | (1.0x) | 1100 | (1.0x) | 13.10% | 1.6x10<sup>-5</sup> |
| 2 | Matched | MVP European | No | 449,042 | (3.7x) | 6252 | (5.7x) | 1.80% | 2x10<sup>-6</sup> |
| 3 | Non-specific | MVP African | Yes | 121,177 | (1.0x) | 1100 | (1.0x) | 6.10% | 9x10<sup>-6</sup> |
| 4 | Non-specific | MVP European | No | 449,042 | (3.7x) | 6252 | (5.7x) | 0.90% | 2.2x10<sup>-5</sup> |

We evaluated the predictive performance (*R*^2^) of ancestry-specific PRISM models in the UKB African ancestry held-out test set across four select traits. To quantify average improvement over the baseline model, we applied orthogonal distance regression (ODR): *R*^2^_PRISM_-*R*^2^_baseline_ ∼ 0+*R*^2^_baseline_ (**Methods**). We show the number of individuals and fine-mapped variants used in each model and the estimated average improvement and statistical significance (p-value) from the ODR regression.

### Tissue-matched annotations enhance PGS transferability

To investigate the impact of tissue specificity in annotations on PGS transferability, we trained a series of PRISM models using tissue-matched, tissue-mismatched, or tissue-non-specific annotations for comparison of their predictive performance (**Methods**), while fixing the fine-mapped variants to those obtained from MVP African source population. We refer to tissue-matched and tissue-mismatched collectively as tissue-specific, as both are restricted to a single tissue. We evaluated the relative improvement in predictive performance compared to the baseline model, the normalized performance relative to the estimated trait heritability (*R^2^/h^2^*), and the average improvement across the four select traits (**Fig. 2**).

**Figure 2.**
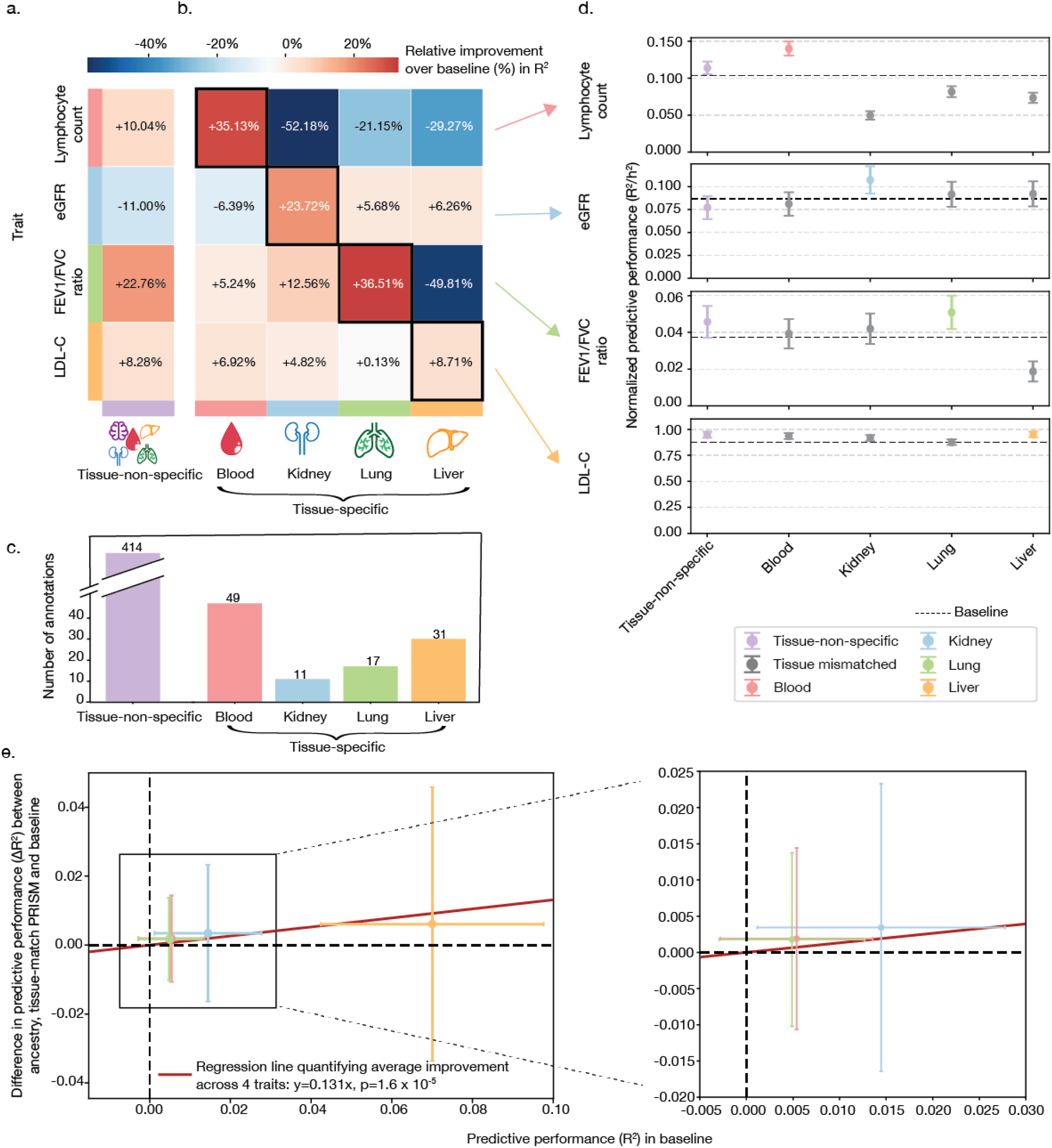
Tissue-matched PRISM improves PGS transferability across four select traits in African ancestry individuals in the UK Biobank. **a, b.** Relative improvements (%) in predictive performance (*R*^2^) over the baseline model for tissue-non-specific (**a**) and tissue-specific (**b**) PRISM models. In (**b)**, we outline tissue-matched PRISM models with black diagonal boxes. **c.** Total number of annotations used in each model, with tissue-specific annotations colored by tissue. **d.** Predictive performance of each PRISM model normalized by estimated trait heritability (*R*^2^/*h*^2^). Colors indicate tissue-specificity. **e.** Average improvement in predictive performance across four traits, quantified using orthogonal distance regression (ODR): *R*^2^_PRISM_-*R*^2^_baseline_ ∼ 0+*R*^2^_baseline_ (**Methods**). Error bars represent the 95% confidence intervals.

Overall, tissue-matched PRISM models showed the greatest improvements in predictive performance (*R*^2^), despite using 8-38 times fewer annotations than tissue-non-specific models (**Fig. 2a-c**). Specifically, using 11-49 annotations, we observed a 36.51% improvements in predictive performance for FEV_1_/FVC ratio (lung; 17 annotations), 35.13% for lymphocyte count (blood; 49 annotations), 23.72% for eGFR (kidney; 11 annotations), and 8.71% for LDL-C (liver; 31 annotations), relative to the baseline model, which does not rely on annotations or fine-mapping results (**Fig. 2b**). In contrast, tissue-non-specific PRISM models trained on the full set of 414 annotations resulted in limited or negative gains: 22.76% for FEV_1_/FVC ratio, 10.04% for lymphocyte count, −11.00% for eGFR, and 8.28% for LDL-C (**Fig. 2a**). We observed the advantage of tissue-matched PRISM across both commonly (e.g., blood with 47 annotations) and less (kidney with only 11 annotations) well-represented tissues, despite substantial variability in annotation availability (**Fig. 2c-d**, **Methods**). Across traits, tissue-matched PRISM led to a 13.1% (95% CI:[3.3%, 23%], p=1.6×10^−5^) average improvement in predictive performance (**Fig. 2e**, **Table 1**). We used a geometric mean of 23.1 annotations for tissue-matched PRISM models, which is a 17.9-fold decrease compared to the tissue-non-specific model. Overall, these results highlight that curated, biologically relevant annotations are more informative than broader, less-specific annotations for enhancing PGS transferability.

### Comparison to existing approaches for PGS transferability

To assess the advantage of PRISM, we compared its predictive performance against those from three widely-used PGS approaches (**Fig. 3a**): IMPACT[15], a representative transcription factor-binding-based approach that relies solely on one type of annotation but with a larger number of 707 spanning different tissues or cell-types; SBayesRC-multi[21], which combines tissue non-specific annotations with multi-ancestry modeling; and PolyPred+[20], which integrates all three main strategies but in a less comprehensive manner than PRISM. Specifically, we focused on the predictive performance (*R^2^*) in the same set of African ancestry individuals in the held-out test set (**Methods**). Overall, PRISM demonstrated competitive predictive performance for all tested traits with three out of four traits with the best predictive performance and second-best for lymphocyte count, where it ranked after SBayesRC-multi (**Fig. 3b**, **Supplementary Table 2**). Together, these results demonstrate that PRISM is a competitive and robust method for enhancing PGS transferability.

**Figure 3.**
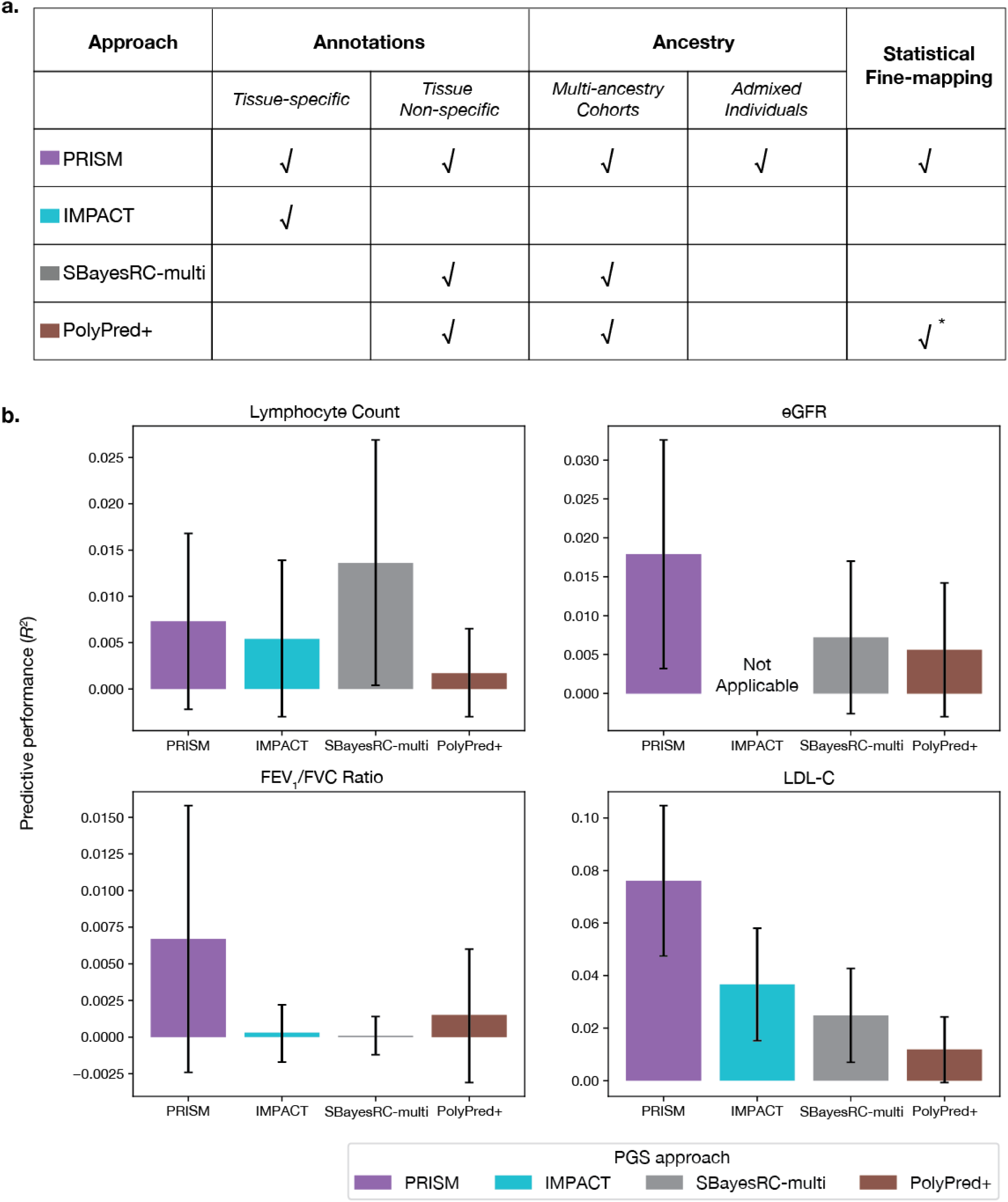
PRISM shows improved and competitive predictive performance for all traits against existing approaches. **a.** Comparison of the PGS approaches given their strategies for enhancing PGS transferability. PRISM is the most comprehensive approach in integrating all three complementary strategies. √ indicates the approach uses the strategy. For fine-mapping, PolyPred+ performs fine-mapping directly (√*), whereas PRISM uses fine-mapping results, offering more flexibility (√). **b.** Predictive performance (*R^2^*) of annotation-informed approaches (color) across selected traits. PRISM shows improved and competitive predictive performance for all traits against existing approaches. Error bars represent 95% confidence intervals. We note that IMPACT is not applicable to eGFR, as no suitable lead annotation was identified (**Methods**).

### PRISM enables biological interpretation of selected variants

Having demonstrated that ancestry- and tissue-matched annotations enhance PGS performance, we next performed biological interpretation by leveraging the sparsity of PRISM models. Specifically, we prioritized variants, examined their contributing annotations, and quantified the importance of annotations (**Fig. 4**). As an illustrative example, we focused on the ancestry- and tissue-matched PRISM model for FEV_1_/FVC ratio which yielded the largest enhancement in PGS transferability, and presented results for the other three traits in **Supplementary Figs. 2-3, 5-7**.

**Figure 4.**
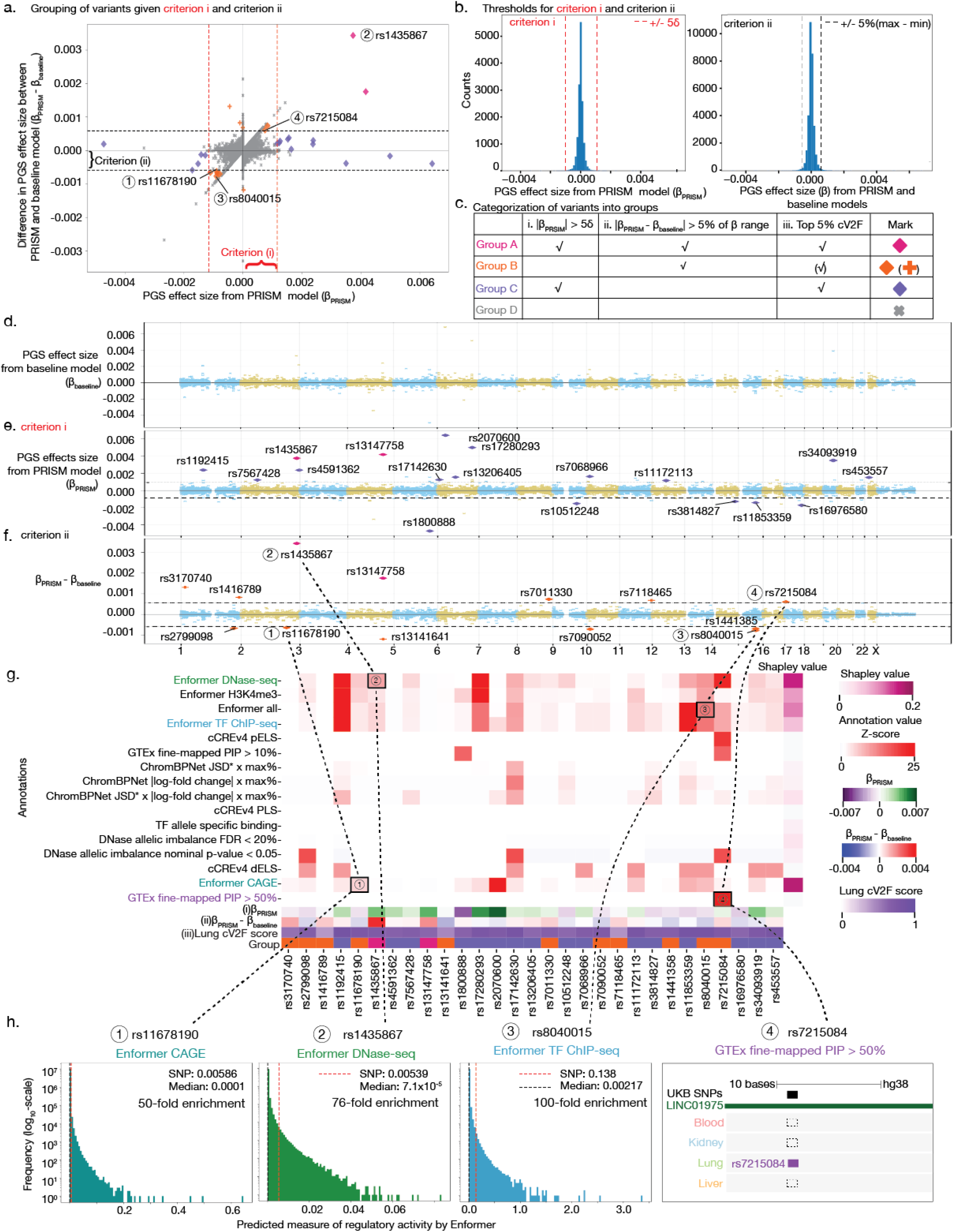
Biological interpretation of ancestry- and tissue-matched PRISM-selected variants for FEV1/FVC ratio using lung-specific annotations. **a.** We use three criteria to group variants (color) (**Methods**). We plot criterion i (absolute value of PRISM effect size) on the x-axis, criterion ii (absolute value of difference between PRISM and baseline effect sizes) on the y-axis, and indicate criterion iii (top 5% cV2F score) with diamond shapes. **b.** We show thresholds used for criteria i and ii. We use five standard deviations for criterion i (left) and the top 5% of the effect size difference range for criterion ii (right). **c.** We assign variants to groups based on a combination of the three criteria. **d-f**. We show genome-wide PGS effect sizes from the baseline model (**d**), PRISM model (**e**), and their differences (**f**). In **e-f**, the blue lines represent thresholds for criteria i and ii. Color and shape indicate group assignment. **g**. We visualize absolute Z-scores of annotation values (y-axis) for prioritized variants (x-axis). We display the three criteria and group assignment at the bottom, along with genome-wide Shapley values on the right. We cap Z-scores at 25 for visualization; the full-scale version is in **Supplementary Fig. 4**. **h**. We show histograms and browser tracks of annotations for four prioritized variants. Abbreviations. JSD: Jensen–Shannon Divergence; TF: transcription factor.

To prioritize genetic variants for interpretation, we applied three criteria: (i) the absolute value of the effect size from the PRISM PGS model (|β_PRISM_|), (ii) the absolute value of the difference from the annotation-agnostic baseline model (|β_PRISM_-β_Baseline_|), and (iii) the continuous cV2F score (**Fig. 4a-b**). Based on these criteria, we defined four variant groups (**Fig. 4c**, **Methods**, **Supplementary Table 3a-d**). Briefly, Group A variants satisfied all three criteria; Group B and Group C satisfied subsets of them; and Group D satisfied none. As expected, most prioritized variants were non-coding and broadly distributed across the genome (**Fig. 4d-f**, **Supplementary Table 4**).

Focusing on the prioritized variants in Groups A-C, we investigated the contributions of annotations (**Fig. 4g**). To quantify the importance of annotations, we used Z-scores to capture local, per-variant contributions and Shapley values to estimate genome-wide, global relevance (**Methods**)[38]. For example, Enformer tracks revealed strong regulatory signals for several prioritized variants[24]: rs11678190 showed a 50-fold higher CAGE value than the genome-wide median (0.0058 vs. 0.0001); rs1435867 exhibited a 76-fold enrichment in DNase-seq predictions (0.00538 vs. 7.1×10^−5^); and rs8040015 revealed a 100-fold increase in transcription factor ChIP-seq signal (0.138 vs. 0.00217) (**Fig. 4h**). In addition, rs7215084 overlapped a tissue-specific fine-mapped quantitative trait locus (eQTL) in the lung[20], further supporting the biological relevance of the variant. Notably, no single annotation dominated the contributions across prioritized variants, highlighting the importance of integrating diverse annotations within the PRISM.

To understand how annotations contribute to selecting variants in LD, we compared the annotations of selected variants to those of nearby variants. We focused on two variants selected by the ancestry- and tissue-matched PRISM model for FEV_1_/FVC ratio, and visualized their annotations using the UCSC Genome Browser[39].

The first variant, rs11853359 (15:71329185:G:A, GRCh38), is a well-characterized regulatory variant in a putative enhancer of *THSD4* and acts as an eQTL for the same gene in the lung[40,41]. Among eight variants in LD (*r^2^*>0.8, MAF>0.2), PRISM assigned the largest effect size to rs11853359 (**Fig. 5a-b, Methods**). In addition to its eQTL signal, rs11853359 overlaps with a distal enhancer-like element[40], exhibits DNase allelic imbalance, and shows high predicted regulatory activity by Enformer (**Fig. 5c**).

**Figure 5.**
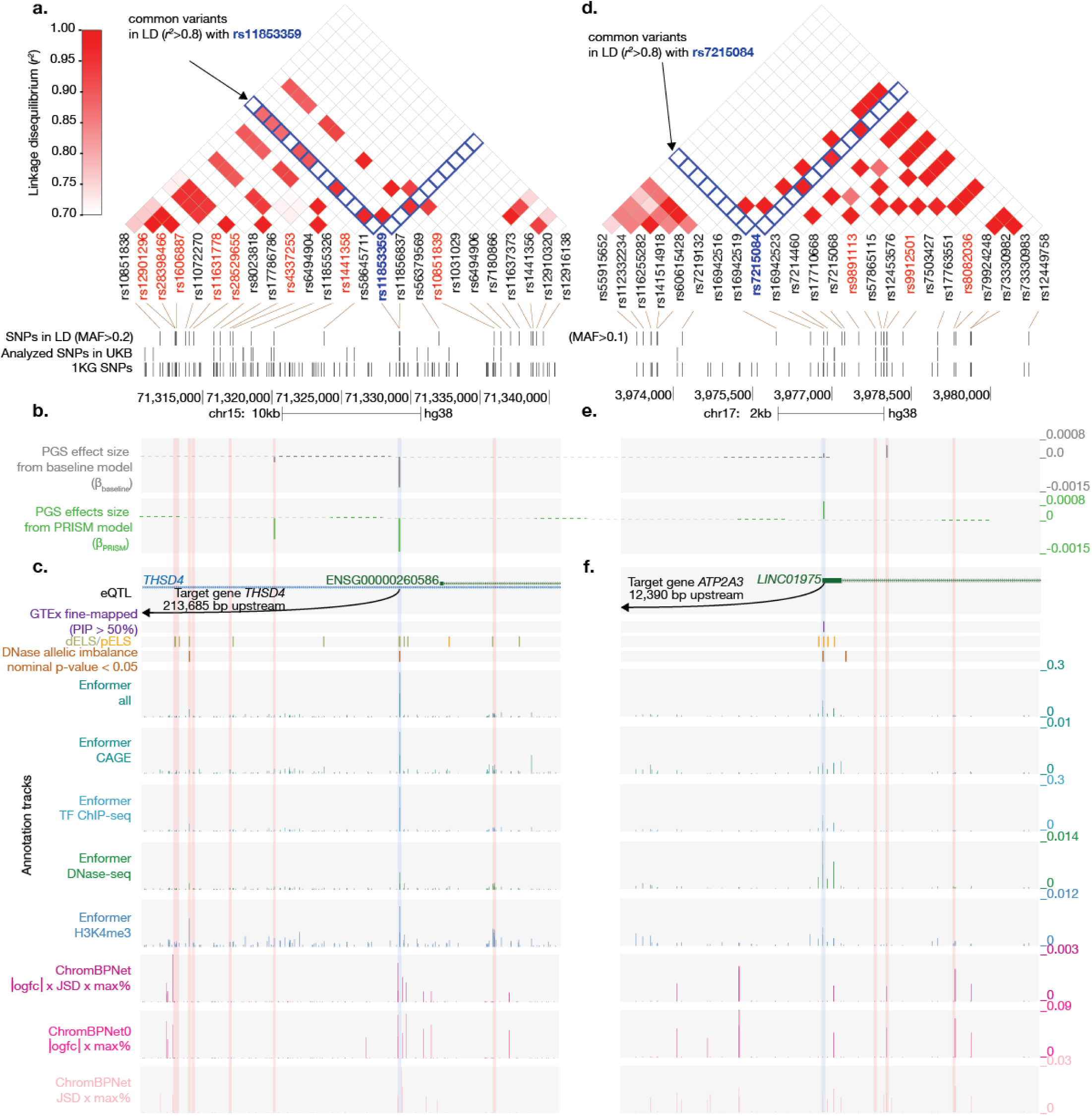
Biological interpretation of PRISM-selected variants for FEV₁/FVC ratio using lung-specific annotations. We highlight two PRISM-selected variants: rs11853359 (**a**–**c**) and rs7215084 (**d**–**f**). For each variant, we show neighboring common variants in linkage disequilibrium (LD; *r*^2^ > 0.8) (**a**, **d**), PGS coefficients from the PRISM model (green) and baseline model (gray) (**b**, **e**), and annotation tracks (**c**, **f**). We highlight the variants of interest in blue and neighboring LD variants in red. **a**–**c**. rs11853359 at the *THSD4* locus. The variant overlaps a distal enhancer-like element (dELS) and acts as an eQTL for *THSD4* expression in the lung. In **a**, we used a MAF threshold of > 0.2 to highlight neighboring variants. **d**–**f**. rs7215084 at the *LINC01975*/*ATP2A3* locus. The variant overlaps proximal enhancer-like elements (pELS) and is a fine-mapped eQTL for *ATP2A3* expression in the lung. In **d**, we used a MAF threshold of > 0.1 to highlight neighboring variants. Abbreviations. 1KG: 1000 Genomes; logfc: log fold change; JSD: Jensen–Shannon divergence; TF: transcription factor.

The second variant, rs7215084 (17:3976854:C:T, GRCh38), is less characterized in the literature, but exhibits stronger annotations than its neighboring variants in LD. Among three variants in LD (*r^2^*>0.8, MAF>0.1), PRISM selected only rs7215084 (**Fig. 5d-e**). Notably, the baseline model selected a different variant in strong LD, rs9912501 (*r^2^*=0.96), which PRISM did not select. A previous study reported rs7215084 as a fine-mapped eQTL for *ATP2A3* in the lung (posterior inclusion probability=89%, p-value:7×10^−27^)[33]. We found that rs7215084 overlaps a proximal enhancer-like element, exhibits DNase allelic imbalance, and shows predicted regulatory activity by Enformer (**Fig. 5f**). Furthermore, *ATP2A3*, the putative regulatory downstream target, plays a role in calcium sequestration and muscle contraction[42,43]. A rare variant burden test also links *ATP2A3* to asthma, reinforcing its functional importance in airway physiology[44]. Additional Hi-C and JASPAR data support a 3D chromatin interaction between rs7215084 and *ATP2A3*, with the variant located in a predicted ZNF454 binding site[45,46].

Overall, these examples demonstrate the ability of PRISM to effectively combine multimodal annotations specific to the trait of interest, prioritizing more biologically relevant genetic variants in PGS models.

## Discussion

We present PRISM, the first systematic approach for improving polygenic score transferability by integrating large-scale genomic annotations with ancestry- and tissue-aware modeling across the continuum of genetic ancestry. PRISM unifies three complementary strategies in a maximally integrative framework: integration of genomic annotations, multi-ancestry modeling, and incorporation of fine-mapping results. Here, we applied PRISM to the most comprehensive integrative effort to date, combining 7352 fine-mapped variant-trait pairs from MVP with 414 annotations from ENCODE[11,22]. Using transfer learning, we first learned the optimal combination of annotations and then utilized it to prioritize variants in PGS training within the UKB[28]. For individuals of African ancestry, we observed the greatest improvements in predictive performance when using ancestry- and tissue-matched PRISM models. PRISM showed competitive and improved predictive performance for existing PGS approaches[15,20,21]. PRISM also facilitates biological interpretation by highlighting selected variants that show clear annotation support for trait relevance.

Biologically, PRISM and its empirical applications to the largest-to-date resources highlights several strategic implications for developing equitable polygenic scores. First, we found that no single annotation would be sufficient to enhance the predictive performance, calling for the need of integrative strategies like ours. Second, in combining multiple annotations, we found that biological alignment can outweigh over 100-fold differences in data availability in terms of genetic ancestry and tissue-specificity. In our empirical analysis, ancestry-matched fine-mapping variants were 5.7 times fewer in African source population than that in the European counterpart and tissue-specific models relied on an 17.9-fold smaller number of annotations. Combined, this corresponds to a 102-fold difference in available annotations. Despite this stark contrast, PRISM models that incorporate biologically relevant annotations demonstrated average improvement of 13.1% over the baseline model. These findings have important implications. On the one hand, they underscore the need to expand data collection across diverse genetic ancestries, biosample types, and environmental contexts to develop more comprehensive and representative annotation resources. On the other hand, they offer a pragmatic approach in the meantime: integrating the most biologically relevant available resources can already yield meaningful benefits in predictive accuracy and equity, even before more comprehensive data become available.

Methodologically, PRISM unifies three complementary strategies to enhance PGS transferability within a maximally integrative framework. First, it enables the incorporation of a variety of annotations, spanning continuous and binary data types, tissue- and cell line-specific biosample coverages, both variant- and element-level features, and sources ranging from experimental assays to large-scale predictive models. These heterogeneous inputs are standardized into annotation scores that guide penalty factor assignment in model training. Second, PRISM supports multi-ancestry training not only across distinct uni-ancestry cohorts but also by incorporating admixed individuals, thereby leveraging additional diversity to improve generalizability. This design makes PRISM broadly applicable across the continuum of genetic ancestry, aligning with the growing recognition that ancestry is not a set of discrete categories[5,21,47]. Third, rather than performing fine-mapping itself, PRISM integrates existing fine-mapping results, providing flexibility to use outputs from different tools while maintaining a consistent framework.

In our empirical comparison to main existing approaches for PGS transferability, PRISM shows improved and competitive predictive performance for all traits. PRISM consistently outperformed IMPACT, which included a larger number of annotations but was restricted to a single type, predicted transcription factor binding. By incorporating a wider spectrum of annotation that captures broader biological context and explicitly modeling ancestry, PRISM achieved stronger enhancement in PGS transferability, illustrating no single annotation is sufficient to capture the underlying biology. Relative to SBayesRC-multi, PRISM delivered comparable predictive performance across traits, with the only exception being LDL-C. This difference is likely due to the underlying annotation resources: SBayesRC-multi leveraged baseline-LD v2.2[48–50], which includes FANTOM5 enhancer annotations previously shown to be strongly enriched for immunological diseases, likely offering more biologically relevant explanatory power for lymphocyte count[51]. Finally, PRISM consistently outperformed PolyPred+, achieving similar annotation-based prioritization without the need for LD reference panels. This is a major advantage when analyzing underrepresented or admixed populations, where appropriate reference panels are often unavailable. Indeed, in our application of PolyPred+, 14.5% genomic regions were excluded from fine-mapping due to the lack of overlapping SNPs across the summary statistics, priors, and LD reference panel. Across all comparisons, PRISM proved more comprehensive than existing approaches and less constrained by data availability.

The modular design of PRISM allows systematic investigation of ancestry effects across multiple stages of PGS development by modeling four key components: (1) the source population for fine-mapping, (2) the biosample ancestry for annotations, (3) the ancestry of individuals used for PGS model development, and (4) the ancestry of the test set used for evaluation (i.e., the target population). In practice, annotations derived from diverse ancestry groups remain limited[26]. However, we showed that integrating existing available annotations with ancestry-specific fine-mapping results still improved PGS transferability, likely reflecting the shared biology across populations. Moreover, our previous work on iPGS demonstrated that the direct inclusion of minority and admixed individuals can substantially improve predictive performance[18]. Overall, these findings provide practical guidance for optimizing PGS in underrepresented populations utilizing available resources.

There are several directions for future studies. First, several implementation choices currently rely on empirically motivated heuristics (e.g., variant grouping thresholds and penalty factor assignment); data-driven optimization of the parameters would be helpful. Second, we analyzed traits with one clearly defined primary tissue; future work should incorporate multi-tissue models to capture complex regulatory architectures, potentially through statistical decomposition, pathway-based partitioning, or pleiotropy-informed analysis[52,53]. Third, identifying the most biologically relevant annotations currently requires manual curation; future approaches could leverage metadata-informed selection strategies, combined with small-scale validation, to enhance the broader applicability. Fourth, our study currently focuses on single-trait PGS; extending PRISM to cross-trait and multi-trait settings may improve performance by leveraging shared genetic architecture, particularly for underpowered traits[54–56]. Fifth, this initial application focuses on individuals of African ancestry in the UKB; future studies should expand the application to additional populations, leveraging resources from All of Us, MVP, and other emerging resources[10–12]. Lastly, PRISM’s modular framework allows integration with other PGS approaches for optimal performance. For instance, the multi-ancestry modeling component could be implemented with PRS-CSx[17] instead of iPGS[18]. Similarly, SNP-level effect-size derivation could be replaced with a Bayesian framework that allows incorporation of multiple priors[57].

Overall, our results highlight the advantage of integrating biologically relevant genomic annotations to enhance PGS transferability. We demonstrate that ancestry- and tissue-aware integration can outweigh the benefits of 100 times larger but less specific annotations. The modular design of PRISM offers a pragmatic strategy for enhancing PGS transferability in underrepresented populations: integrating a potentially smaller amount of the most biologically relevant curated resources can offer immediate benefits while waiting for more comprehensive data collection from diverse populations. We make the coefficients of the PGS models available at the ENCODE portal (https://www.encodeproject.org/), the PGS catalog, and the iPGS Browser (https://ipgs.mit.edu/).

## Methods

### Compliance with ethical regulations and informed consent

This research has been conducted using the UK Biobank Resource under Application Number 21942, “Integrated models of complex traits in the UK Biobank” (https://www.ukbiobank.ac.uk/enable-your-research/approved-research/integrated-models-of-complex-traits-in-the-uk-biobank). All participants of UK Biobank provided written informed consent (more information is available at https://www.ukbiobank.ac.uk/explore-your-participation/basis-of-your-participation/).

### The study population in UK Biobank

UK Biobank is a population-based cohort study with genomic and phenotypic datasets across about 500,000 volunteers collected across multiple sites in the United Kingdom[27,28]. We focused on N=406,659 unrelated individuals with genetic data based on the following quality control criteria: (1) used to compute principal components (UKB Data Field 22020); (2) removal of sex mismatch between the sex field in the genotype dataset and phenotype sex (Data Field 31); (3) not reported in “outliers for heterozygosity or missing rate” (Data Field 22027); (4) not reported in “sex chromosome aneuploidy” (Data Field 22019); and (5) do not have ten or more third-degree relatives (Data Field 22021)[53,56,58]. We used a combination of genetic principal components (Data Field 22009) and self-reported ethnic background (Data Field 21000) to define four population groups: white British (WB), non-British white (NBW), African (Afr), and South Asian (SA), and kept the remaining unrelated individuals as Others (**Supplementary Table 1**)[56]. We used the same training set (70%) for PGS model fitting, validation set (10%) for determining sparsity of PGS models, and test set (20%) for evaluating predictive performance, as described in the previous study[18].

### Variant annotation and quality control in UK Biobank

For the UKB resource, we used the directly genotyped dataset (release version 2), imputed genotypes (release version 3), imputed HLA allelotype (release version 2), and GRCh37 human reference genome[27]. We used variant annotation with Ensembl’s Variant Effect Predictor (VEP) (version 101) with the LOFTEE plugin and ClinVar[18,59–62] as in our previous study. We grouped the VEP-predicted consequence of the variants into six groups: protein-truncating variants (PTVs), protein-altering variants (PAVs), proximal coding variants (PCVs), intronic variants (Intronic), genetic variants on untranslated regions (UTR), and other non-coding variants (Others)[63]. We considered “pathogenic” and “likely pathogenic” variant annotations from ClinVar[64].

We used the same variant-level quality control criteria as in our previous study[18]. For the directly genotyped dataset, we focused on variants passing the following criteria: (1) the missingness of the variant is less than 1% given the two genotyping arrays (the UK BiLEVE Axiom array and UKB Axiom array) cover a slightly different set of variants and (2) the Hardy-Weinberg disequilibrium test p-value greater than 1.0×10^−7^. For the imputed genotype dataset, we used the following criteria: (1) the missingness of the variant is less than 1%; (2) minor allele frequency (MAF) greater than 0.01%; (3) imputation quality score (INFO score) greater than 0.3; (4) is not present in the directly genotyped dataset; and (5) present in the HapMap Phase 3 dataset[18]. For the HLA allelotype, we kept the imputed allelotype dosage within [0, 0.1), (0.9, 1.1), or (1.9, 2.0] and converted it to hard calls[27,65]. We focused on the HLA allelotype with (1) missingness no more than 1% and (2) Hardy-Weinberg disequilibrium test p-value greater than 1.0×10^−4^. We concatenated all variants and allelotypes into one dataset using PLINK 2.0 (v2.00a3.3LM 3 Jun 2022)[66]. This resulted in a total of 1,316,181 variants considered in the analysis.

### Phenotype definition

We focused on four select traits with clear primary tissue to test the utility of tissue-specific annotations in polygenic prediction: blood-lymphocyte count, kidney-eGFR, liver-LDL-C, and lung-FEV_1_/FVC ratio[36]. We used the race-neutral CKD-EPI (Creatinine-Cystatin C) equation to define eGFR values[67]. Those phenotypes were collected in up to 3 instances: (1) the initial assessment visit (2006–2010), (2) the first repeat assessment visit (2012–2013), and (3) the imaging visit (2014–present). For each individual, we took the median of non-missing values. We show the number of individuals for the four traits in **Supplementary Table 1**.

### Genome-wide association analysis and heritability estimation

We conducted genome-wide association analysis (GWAS) using PLINK (version 2.00 alpha) as in our previous study[18,66]. Briefly, we used population-specific genotype principal components (PCs) characterized with the randomized algorithm to account for population structure[68]. We included the top 10 genotype PC loadings as well as age, sex, Townsend deprivation index, and genotyping array as covariates using the plink2 command “--glm zs omit-ref no-x-sex log10 hide-covar skip-invalid-pheno cc-residualize firth-fallback”[69]. Approximately 10% of UKB participants were genotyped using the UK BiLEVE Axiom array, while the rest were genotyped using the UKB Axiom array[27]. For genetic variants directly measured on both arrays, we included an “array” indicator variable as a covariate and specified whether the UK BiLEVE Axiom array or the UKB Axiom array was used for genotyping. We conducted separate GWAS for UKB WB and Afr individuals, with training sample sizes varying depending on the specific PGS approach applied (see **Methods**: Application of existing approaches for PGS transferability). We applied GWAS results estimated in WB individuals to compute SNP-based heritability for each trait, using linkage disequilibrium score regression (LDSC) with 1000 Genomes Phase 3 European-ancestry individuals as the LD reference [6,70].

### Generating continuous cV2F scores

We curated fine-mapped variants across 936 traits within the MVP cohort, including 1100 fine-mapped variants from 121,177 individuals of African ancestry and 6552 fine-mapped variants from 449,042 individuals of European ancestry[11]. We curated a total of 414 annotations, of which 272 are tissue-specific (**Supplementary Table 5, Data and code availability**)[11,22,29–33]. We focused on four tissue categories for tissue-specific annotation: blood (derived from blood tissue and the K562 and GM12878 cell lines), kidney, lung, and liver (including annotations from liver tissue and the HepG2 cell line).

To aggregate annotations into variant-level numeric scores, we applied a gradient-boosting model to fine-mapped variants and annotations[34]. The model uses fine-mapped variants with posterior inclusion probability (PIP) > 90% as positive and PIP <1% as negative labels using leave-one-chromosome-out cross-validation. It trains on annotations associated with the GWAS fine-mapped variants, resulting in a continuous cV2F score ranging from 0 to 1[34]. We scored 9,991,229 variants (minor allele count ≥5) genotyped in 1000 Genomes Europeans using the trained models. We repeated the analysis 10 times, corresponding to the combination of the two source ancestries of fine-mapped results (European and African in MVP) and tissue-specificity of annotations, i.e., all (tissue-non-specific), blood, kidney, lung, and liver (**Supplementary Table 6**). We used all 414 annotations for tissue-non-specific cV2F models, whereas we used a subset of annotations in tissue-specific cV2F scores (**Supplementary Table 7**).

### Baseline and PRISM model training

To assess the impact of incorporating annotations and fine-mapping results into PGS training, we fit a baseline and PRISM models and compare their predictive performance. We trained PGS on individual-level genetic data in the UKB. Specifically, we used the iPGS approach, a penalized regression directly learned on the individual-level data that minimizes the following loss function[18]:

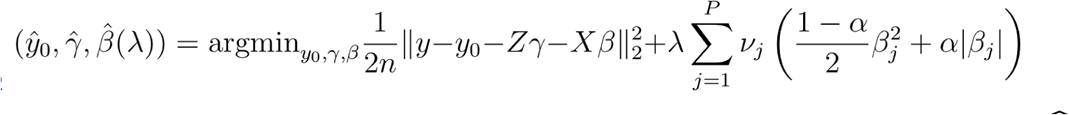

where *y* is the phenotype, *Z* is the covariates, and *X* is the genetic predictors. Here, 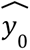 is the estimated intercept of the regression model, and γ̂, and β̂ are the estimated coefficients for covariates and genetic predictors, respectively. We control the sparsity of the solution via the tuning parameter λ, which is optimized on the basis of the predictive performance of the validation set. Covariates include age, sex, Townsend deprivation index, and top 18 genotype PC loadings. We set the elastic net parameter α to 0.99. We grouped variants according to their predicted consequences and genomic annotations and assigned a penalty factor ν_*j*_ to each group. The baseline and PRISM models differ in their penalty factor assignments as described below, based on their biological relevance to our trait of interest. The PRISM models varied by ancestry and tissue specificity, given the type of cV2F score used, resulting in a total of 10 models for each trait (**Supplementary Table 8**).

We used a heuristic to assign penalty factors(**Supplementary Table 9**). For the baseline model, we assigned penalty factors based solely on predicted consequences. For PRISM models, we additionally considered aggregated annotations to assign penalty factors. We set penalty factor cutoffs differently based on ancestry-specific cV2F scores. For African cV2F scores, we binned the values and assigned lower penalty factors to variants falling within the top 5% bin. For European cV2F scores, we set the cutoff to match the number of variants within the top 5% of the African cV2F scores. The full list of cutoffs can be found in **Supplementary Table 6.** To ensure consistency across datasets, we used rsIDs to identify and match UKB variants with their assigned continuous cV2F scores. We matched 1,188,579 out of a total of 1,316,181 variants. Variants without a continuous cV2F score will be assigned a default penalty factor of 1 unless they meet the criteria based on predicted consequences.

For LDL-C, we excluded variants within the *APOE* region due to the known strong genetic influence at this locus and recent proposals of separately modeling polygenic background and strong-acting alleles as in the case for PGS for Alzheimer’s disease[70–74]. Specifically, we excluded rs7412 and nearby variants in LD (*r*^2^>.1) (684 variants in chr19:45,176,340-45,447,221). We evaluated PGS predictive performance against non-*APOE* heritability of the trait[70].

### Evaluating PGS model performance

We evaluated the predictive performance (*R^2^*) of PRISM and baseline models in African ancestry individuals in the UKB (n=1154 for lymphocyte count, n=1134 for eGFR, n=1078 for FEV_1_/FVC ratio, and n=1130 for LDL-C) for: (1) genotype-only models, (2) covariate-only models, and (3) full models combining covariates and genotypes. We reported the predictive performance of genotype-only models in the remainder of the main text unless indicated otherwise.

To quantify the relative improvement of PRISM model predictive performance for each trait, we compared the ancestry- and tissue-matched, the tissue-non-specific, and tissue-mismatched PRISM models each to the baseline model, reporting the percentage increase in *R*^2^ and *R*^2^ normalized by estimated trait heritability.

To quantify the average improvement of the predictive performance of PRISM models over that of the baseline model, we applied ODR, in the form of *R*^2^_PRISM_-*R*^2^_Baseline_ ∼ 0+*R*^2^_Baseline_, where the regression model considers the uncertainties on both variables across the four traits. Specifically, we quantified average improvements for 4 PRISM models: (1) ancestry-mismatched, tissue-non-specific PRISM vs baseline, (2) ancestry-matched, tissue-non-specific PRISM vs baseline, (3) ancestry-mismatched, tissue-matched PRISM vs baseline, and (4) ancestry-matched, tissue-matched PRISM vs baseline.

### Application of existing approaches for PGS transferability

To evaluate the advantage of PRISM in improving predictive performance, we compared PRISM against three widely used approaches: PolyPred+[20], SBayesRC-multi[21], and IMPACT[15]. Across all methods, we evaluated the performance (*R*^2^) in the same held-out set of African ancestry individuals in UKB (n=1154 for lymphocyte count, n=1134 for eGFR, n=1078 for FEV_1_/FVC ratio, and n=1130 for LDL-C). We outline implementation details below.

#### PolyPred+

We ran fine-mapping in unrelated UKB WB individuals (n=270,920) using precomputed priors and an LD reference panel from UKB participants of European ancestry, both provided by the authors via their GitHub repository (https://github.com/omerwe/polyfun)[20]. We restricted our analysis to SNPs (n=1,232,022) present in our analysis cohort with both prior probabilities and LD reference data available. During fine-mapping, up to 14.5% of genomic regions were excluded due to having no overlapping SNPs across summary statistics, priors, and LD reference. In parallel, we estimated tagging effect sizes in UKB Afr individuals (n=4853). Subsequently, we linearly combined them with our fine-mapped effect sizes to construct cross-ancestry PGS for downstream evaluation. We excluded SNPs that lie within the *APOE* region when constructing PGS for LDL-C.

#### SBayesRC-multi

We first applied single-ancestry SBayesRC on UKB WB (n=270,920) and Afr (n=4246) individuals separately, using annotations and LD reference panel from UKB participants of European ancestry, both provided by the authors via their GitHub repository (https://github.com/zhilizheng/SBayesRC)[21]. The resulting SNP effect sizes were used to construct two separate PGS, based on WB and Afr discovery samples, for a separate Afr validation cohort (n=607). We subsequently considered the optimal linear combination of the two using the same validation set individuals, resulting in a SBayesRC-multi model. We evaluate the predictive performance of the model in the held-out test set of individuals of African ancestry. Because the final step combines polygenic scores rather than SNP-level effect sizes, we excluded SNPs within the *APOE* region when constructing PGS for the validation cohort.

#### IMPACT

We identified the lead annotation for each trait by using the closest trait match reported in Supplementary Table 9 of the IMPACT paper[15]. The specific matched pairs were: lymphocyte count with Lympho, FEV1/FVC ratio with FEV1 FVC Smoke, and LDL-C with LDL. No suitable annotation was found for eGFR. We note that the trait matches are approximate rather than exact, reflecting the closest available counterparts reported. We ran GWAS on UKB WB individuals (n = 270,920) as described above and retained SNPs within the top 5% of lead annotation scores for each trait. We then applied LD clumping in Afr individuals (n=6066) to remove variants in LD with r^2^ > 0.2 with a significance threshold for index SNPs of P = 0.5. We performed thresholding in Afr individuals (n=4853) across a range of p-value cutoffs (0.1, 0.03, 0.01, 0.003, 0.001, 3×10^−4^, 1×10^−4^, 3×10^−5^, and 1×10^−5^) to identify the optimal PGS, which was then applied to the evaluation cohort.

### Biological interpretation

For biological interpretation, we selected genomic loci and examined associated annotations in tissue- and ancestry-matched PRISM models that may contribute to improved PGS transferability. To select for genomic locus, we propose three criteria: (i. relative effect size) we require the absolute effect size from the tissue- and ancestry-specific PRISM (β_PRISM_) deviates is greater than five standard deviations, (ii. effect size difference) the absolute value of effect size differences between the baseline and PRISM model (|β_PRISM_-β_Baseline_|) is greater than 5% of the range of the effect sizes from both models, and (iii. variant prioritization) the variant is prioritized based on its continuous AFR cV2F score ranking in the top 5%. We grouped variants given the combination of these three criteria: Group A variants fulfill all three criteria, Group B variants fulfill either criterion ii or iii, and Group C variants fulfill criterion i. Group D variants fulfill none of the criteria (**Fig. 4b-c**).

To examine each annotation’s contribution to variant selection in the ancestry- and tissue-matched PRISM model, we quantified their genome-wide and local importance. We used Shapley value to measure the genome-wide importance of each annotation when generating the continuous cV2F scores[34,38]. For local importance, we calculated the Z-score for the annotation value of each variant. The higher the Shapley value and the absolute Z-score, the greater the annotation’s influence on variant selection.

To visualize annotations in genomic contexts, we focused on FEV_1_/FVC ratio, a lung capacity measurement, as ancestry- and tissue-match PRISM showed the highest improvement in predictive performance (36.51%) for this trait. We examined example loci surrounding rs11853359 and rs7215084, using the UCSC Genome Browser[39]. We defined the visualization windows for rs11853359 and rs7215084 separately, based on LD and MAF thresholds: (*r^2^*>0.8, MAF>0.2) for rs11853359 and (*r^2^*>0.8, MAF>0.1) for rs7215084. For both LD and MAF, we used the European population from the 1000 Genomes Phase 3 reference panel and applied the following PLINK command: plink --r2 --ld-window 100000000 --ld-window-kb 1000 --ld-window-r2 0.1[6][75]. We showed variants in LD (*r^2^* > 0.1) with the index variant. We showed PGS coefficients for the ancestry- and tissue-matched PRISM model and the baseline model. We loaded annotation tracks only if signals were present for variants within the defined genomic window. For GTEx fine-mapping tracks, we showed the binary indicator track showing variants with PIP>50%[21].

## Acknowledgments

This work was supported in part by the National Institutes of Health grants AG054012, AG058002, MH109978, AG062377, AG081017, NS129032, AG077227, NS110453, NS115064, AG062335, AG074003, NS127187, AG067151, MH119509, HG008155, and DA053631 (M.K.). We thank Jacob C. Ulirsch, Amy Grayson, Patricia Purcell, Martin Wohlwend, and the members of the Kellis lab for their scientific suggestions. We thank Shaila A. Musharoff and Andrew G. Clark for their generous support during the completion of this project. The content is solely the responsibility of the authors and does not necessarily represent the official views of the funding agencies; funders had no role in study design, data collection, and analysis, the decision to publish, or the preparation of the manuscript.

## Author contributions

Y.T. conceived, designed, and supervised the study; X.T. performed data analysis and prepared the figures; X.T. and Y.T. conducted the polygenic score analyses; T.F. and K.K.D. performed the cV2F score analysis; W.F.L. contributed to the biological interpretation of the results; Y.T. and X.T. drafted the manuscript with input from M.K.; M.K. was responsible for funding acquisition; and all authors reviewed and approved the final version of the manuscript.

## Data and code availability

The PRISM pipeline and corresponding analysis are available at: https://github.com/lucy-tian/PRISM/tree/main.

The code to replicate continuous cV2F score is available at: https://github.com/Deylab999MSKCC/cv2f/tree/main.

The BASIL algorithm implemented in the R snpnet package version 2 (https://github.com/rivas-lab/snpnet/tree/compact) was used in the PGS analysis.

We will make resources available at the ENCODE portal (https://www.encodeproject.org/) the PGS catalog, and the iPGS Browser (https://ipgs.mit.edu/) upon acceptance of the manuscript.

ENCODE (Phase 4) datasets, including ChromBPNet scores (Kundaje lab), MPRA allelic skew effects and deep learning model annotations (Tewhey lab), DNase Allelic imbalance calls (Viertsra lab),and cCRE (v4) maps (Moore, Weng labs) are currently available via request to the authors and will be made publicly available on the ENCODE portal.

## References

1. Lewis CM, Vassos E. Polygenic risk scores: from research tools to clinical instruments. Genome Med. 2020;12: 44. doi:10.1186/s13073-020-00742-5

2. Wand H, Lambert SA, Tamburro C, Iacocca MA, O’Sullivan JW, Sillari C, et al. Improving reporting standards for polygenic scores in risk prediction studies. Nature. 2021;591: 211–219. doi:10.1038/s41586-021-03243-6

3. Martin AR, Kanai M, Kamatani Y, Okada Y, Neale BM, Daly MJ. Clinical use of current polygenic risk scores may exacerbate health disparities. Nat Genet. 2019;51: 584–591. doi:10.1038/s41588-019-0379-x

4. Kachuri L, Chatterjee N, Hirbo J, Schaid DJ, Martin I, Kullo IJ, et al. Principles and methods for transferring polygenic risk scores across global populations. Nat Rev Genet. 2023;25: 8–25. doi:10.1038/s41576-023-00637-2

5. Ding Y, Hou K, Xu Z, Pimplaskar A, Petter E, Boulier K, et al. Polygenic scoring accuracy varies across the genetic ancestry continuum. Nature. 2023;618: 774–781. doi:10.1038/s41586-023-06079-4

6. 1000 Genomes Project Consortium. A global reference for human genetic variation. Nature. 2015;526: 68–74. doi:10.1038/nature15393

7. Majara L, Kalungi A, Koen N, Tsuo K, Wang Y, Gupta R, et al. Low and differential polygenic score generalizability among African populations due largely to genetic diversity. HGG Adv. 2023;4: 100184. doi:10.1016/j.xhgg.2023.100184

8. Morales J, Welter D, Bowler EH, Cerezo M, Harris LW, McMahon AC, et al. A standardized framework for representation of ancestry data in genomics studies, with application to the NHGRI-EBI GWAS Catalog. Genome Biol. 2018;19: 21. doi:10.1186/s13059-018-1396-2

9. Genovese G, Friedman DJ, Ross MD, Lecordier L, Uzureau P, Freedman BI, et al. Association of trypanolytic ApoL1 variants with kidney disease in African Americans. Science. 2010;329: 841–845. doi:10.1126/science.1193032

10. All of Us Research Program Genomics Investigators. Genomic data in the All of Us Research Program. Nature. 2024;627: 340–346. doi:10.1038/s41586-023-06957-x

11. Verma A, Huffman JE, Rodriguez A, Conery M, Liu M, Ho Y-L, et al. Diversity and scale: Genetic architecture of 2068 traits in the VA Million Veteran Program. Science. 2024;385: eadj1182. doi:10.1126/science.adj1182

12. Graham SE, Clarke SL, Wu K-HH, Kanoni S, Zajac GJM, Ramdas S, et al. The power of genetic diversity in genome-wide association studies of lipids. Nature. 2021;600: 675–679. doi:10.1038/s41586-021-04064-3

13. Mulder N, Abimiku A ’le, Adebamowo SN, de Vries J, Matimba A, Olowoyo P, et al. H3Africa: current perspectives. Pharmgenomics Pers Med. 2018;11: 59–66. doi:10.2147/PGPM.S141546

14. Kullo IJ, Conomos MP, Nelson SC, Adebamowo SN, Choudhury A, Conti D, et al. The PRIMED Consortium: Reducing disparities in polygenic risk assessment. Am J Hum Genet. 2024;111: 2594–2606. doi:10.1016/j.ajhg.2024.10.010

15. Amariuta T, Ishigaki K, Sugishita H, Ohta T, Koido M, Dey KK, et al. Improving the trans-ancestry portability of polygenic risk scores by prioritizing variants in predicted cell-type-specific regulatory elements. Nat Genet. 2020;52: 1346–1354. doi:10.1038/s41588-020-00740-8

16. Crone B, Boyle AP. Enhancing portability of trans-ancestral polygenic risk scores through tissue-specific functional genomic data integration. PLoS Genet. 2024;20: e1011356. doi:10.1371/journal.pgen.1011356

17. Ruan Y, Lin Y-F, Feng Y-CA, Chen C-Y, Lam M, Guo Z, et al. Improving polygenic prediction in ancestrally diverse populations. Nat Genet. 2022;54: 573–580. doi:10.1038/s41588-022-01054-7

18. Tanigawa Y, Kellis M. Power of inclusion: Enhancing polygenic prediction with admixed individuals. Am J Hum Genet. 2023;110: 1888–1902. doi:10.1016/j.ajhg.2023.09.013

19. Schaid DJ, Chen W, Larson NB. From genome-wide associations to candidate causal variants by statistical fine-mapping. Nat Rev Genet. 2018;19: 491–504. doi:10.1038/s41576-018-0016-z

20. Weissbrod O, Kanai M, Shi H, Gazal S, Peyrot WJ, Khera AV, et al. Leveraging fine-mapping and multipopulation training data to improve cross-population polygenic risk scores. Nat Genet. 2022;54: 450–458. doi:10.1038/s41588-022-01036-9

21. Zheng Z, Liu S, Sidorenko J, Wang Y, Lin T, Yengo L, et al. Leveraging functional genomic annotations and genome coverage to improve polygenic prediction of complex traits within and between ancestries. Nat Genet. 2024;56: 767–777. doi:10.1038/s41588-024-01704-y

22. ENCODE Project Consortium, Moore JE, Purcaro MJ, Pratt HE, Epstein CB, Shoresh N, et al. Expanded encyclopaedias of DNA elements in the human and mouse genomes. Nature. 2020;583: 699–710. doi:10.1038/s41586-020-2493-4

23. GTEx Consortium. The Genotype-Tissue Expression (GTEx) project. Nat Genet. 2013;45: 580–585. doi:10.1038/ng.2653

24. Avsec Ž, Agarwal V, Visentin D, Ledsam JR, Grabska-Barwinska A, Taylor KR, et al. Effective gene expression prediction from sequence by integrating long-range interactions. Nat Methods. 2021;18: 1196–1203. doi:10.1038/s41592-021-01252-x

25. Pampari A, Shcherbina A, Kvon EZ, Kosicki M, Nair S, Kundu S, et al. ChromBPNet: bias factorized, base-resolution deep learning models of chromatin accessibility reveal cis-regulatory sequence syntax, transcription factor footprints and regulatory variants. bioRxiv. 2025. doi:10.1101/2024.12.25.630221

26. Breeze CE, Beck S, Berndt SI, Franceschini N. The missing diversity in human epigenomic studies. Nat Genet. 2022;54: 737–739. doi:10.1038/s41588-022-01081-4

27. Bycroft C, Freeman C, Petkova D, Band G, Elliott LT, Sharp K, et al. The UK Biobank resource with deep phenotyping and genomic data. Nature. 2018;562: 203–209. doi:10.1038/s41586-018-0579-z

28. Sudlow C, Gallacher J, Allen N, Beral V, Burton P, Danesh J, et al. UK biobank: an open access resource for identifying the causes of a wide range of complex diseases of middle and old age. PLoS Med. 2015;12: e1001779. doi:10.1371/journal.pmed.1001779

29. Abramov S, Boytsov A, Bykova D, Penzar DD, Yevshin I, Kolmykov SK, et al. Landscape of allele-specific transcription factor binding in the human genome. Nat Commun. 2021;12: 2751. doi:10.1038/s41467-021-23007-0

30. Siraj L, Castro RI, Dewey H, Kales S, Nguyen TTL, Kanai M, et al. Functional dissection of complex and molecular trait variants at single nucleotide resolution. bioRxiv. 2024. doi:10.1101/2024.05.05.592437

31. Brennan KJ, Weilert M, Krueger S, Pampari A, Liu H-Y, Yang AWH, et al. Chromatin accessibility in the Drosophila embryo is determined by transcription factor pioneering and enhancer activation. Dev Cell. 2023;58: 1898–1916.e9. doi:10.1016/j.devcel.2023.07.007

32. Nair S, Ameen M, Sundaram L, Pampari A, Schreiber J, Balsubramani A, et al. Transcription factor stoichiometry, motif affinity and syntax regulate single-cell chromatin dynamics during fibroblast reprogramming to pluripotency. bioRxiv. 2023. doi:10.1101/2023.10.04.560808

33. Wang QS, Kelley DR, Ulirsch J, Kanai M, Sadhuka S, Cui R, et al. Leveraging supervised learning for functionally informed fine-mapping of cis-eQTLs identifies an additional 20,913 putative causal eQTLs. Nat Commun. 2021;12: 3394. doi:10.1038/s41467-021-23134-8

34. Fabiha T, Evergreen I, Kundu S, Pampari A, Abramov S, Boytsov A, et al. A consensus variant-to-function score to functionally prioritize variants for disease. bioRxiv. 2024. doi:10.1101/2024.11.07.622307

35. Aw AJ, McRae J, Rahmani E, Song YS. Highly parameterized polygenic scores tend to overfit to population stratification via random effects. bioRxiv. 2024. doi:10.1101/2024.01.27.577589

36. Kanai M, Akiyama M, Takahashi A, Matoba N, Momozawa Y, Ikeda M, et al. Genetic analysis of quantitative traits in the Japanese population links cell types to complex human diseases. Nat Genet. 2018;50: 390–400. doi:10.1038/s41588-018-0047-6

37. Virtanen P, Gommers R, Oliphant TE, Haberland M, Reddy T, Cournapeau D, et al. SciPy 1.0: fundamental algorithms for scientific computing in Python. Nat Methods. 2020;17: 261–272. doi:10.1038/s41592-019-0686-2

38. Lundberg SM, Lee S-I. A unified approach to interpreting model predictions. Neural Inf Process Syst. 2017; 4765–4774. doi:10.5555/3295222.3295230

39. Perez G, Barber GP, Benet-Pages A, Casper J, Clawson H, Diekhans M, et al. The UCSC Genome Browser database: 2025 update. Nucleic Acids Res. 2025;53: D1243–D1249. doi:10.1093/nar/gkae974

40. Yao T-C, Du G, Han L, Sun Y, Hu D, Yang JJ, et al. Genome-wide association study of lung function phenotypes in a founder population. J Allergy Clin Immunol. 2014;133: 248–55.e1–10. doi:10.1016/j.jaci.2013.06.018

41. ENCODE Project Consortium. An integrated encyclopedia of DNA elements in the human genome. Nature. 2012;489: 57–74. doi:10.1038/nature11247

42. Korosec B, Glavac D, Volavsek M, Ravnik-Glavac M. ATP2A3 gene is involved in cancer susceptibility. Cancer Genet Cytogenet. 2009;188: 88–94. doi:10.1016/j.cancergencyto.2008.10.007

43. Pruitt KD, Tatusova T, Klimke W, Maglott DR. NCBI Reference Sequences: current status, policy and new initiatives. Nucleic Acids Res. 2009;37: D32–6. doi:10.1093/nar/gkn721

44. Karczewski KJ, Solomonson M, Chao KR, Goodrich JK, Tiao G, Lu W, et al. Systematic single-variant and gene-based association testing of thousands of phenotypes in 394,841 UK Biobank exomes. Cell Genom. 2022;2: 100168. doi:10.1016/j.xgen.2022.100168

45. Song T, Yao M, Yang Y, Liu Z, Zhang L, Li W. Integrative identification by Hi-C revealed distinct advanced structural variations in lung adenocarcinoma tissue. Phenomics. 2023;3: 390–407. doi:10.1007/s43657-023-00103-3

46. Rauluseviciute I, Riudavets-Puig R, Blanc-Mathieu R, Castro-Mondragon JA, Ferenc K, Kumar V, et al. JASPAR 2024: 20th anniversary of the open-access database of transcription factor binding profiles. Nucleic Acids Res. 2024;52: D174–D182. doi:10.1093/nar/gkad1059

47. Hou K, Xu Z, Ding Y, Mandla R, Shi Z, Boulier K, et al. Calibrated prediction intervals for polygenic scores across diverse contexts. Nat Genet. 2024;56: 1386–1396. doi:10.1038/s41588-024-01792-w

48. Gazal S, Marquez-Luna C, Finucane HK, Price AL. Reconciling S-LDSC and LDAK functional enrichment estimates. Nat Genet. 2019;51: 1202–1204. doi:10.1038/s41588-019-0464-1

49. Gazal S, Loh P-R, Finucane HK, Ganna A, Schoech A, Sunyaev S, et al. Functional architecture of low-frequency variants highlights strength of negative selection across coding and non-coding annotations. Nat Genet. 2018;50: 1600–1607. doi:10.1038/s41588-018-0231-8

50. Gazal S, Finucane HK, Furlotte NA, Loh P-R, Palamara PF, Liu X, et al. Linkage disequilibrium-dependent architecture of human complex traits shows action of negative selection. Nat Genet. 2017;49: 1421–1427. doi:10.1038/ng.3954

51. Finucane HK, Bulik-Sullivan B, Gusev A, Trynka G, Reshef Y, Loh P-R, et al. Partitioning heritability by functional annotation using genome-wide association summary statistics. Nat Genet. 2015;47: 1228–1235. doi:10.1038/ng.3404

52. Qi G, Chhetri SB, Ray D, Dutta D, Battle A, Bhattacharjee S, et al. Genome-wide large-scale multi-trait analysis characterizes global patterns of pleiotropy and unique trait-specific variants. Nat Commun. 2024;15: 6985. doi:10.1038/s41467-024-51075-5

53. Tanigawa Y, Li J, Justesen JM, Horn H, Aguirre M, DeBoever C, et al. Components of genetic associations across 2,138 phenotypes in the UK Biobank highlight adipocyte biology. Nat Commun. 2019;10: 4064. doi:10.1038/s41467-019-11953-9

54. Qian J, Tanigawa Y, Li R, Tibshirani R, Rivas MA, Hastie T. Large-scale multivariate sparse regression with applications to UK Biobank. Ann Appl Stat. 2022;16: 1891–1918. doi:10.1214/21-aoas1575

55. Li R, Tanigawa Y, Justesen JM, Taylor J, Hastie T, Tibshirani R, et al. Survival Analysis on Rare Events Using Group-Regularized Multi-Response Cox Regression. Bioinformatics. 2021;37: 4437–4443. doi:10.1093/bioinformatics/btab095

56. Sinnott-Armstrong N, Tanigawa Y, Amar D, Mars N, Benner C, Aguirre M, et al. Genetics of 35 blood and urine biomarkers in the UK Biobank. Nat Genet. 2021;53: 185–194. doi:10.1038/s41588-020-00757-z

57. Wu H, Pérez-Rodríguez P, Boehnke M, Cui Y, Liang X, Vazquez AI, et al. Improving polygenic score prediction for underrepresented groups through transfer Learning. medRxiv. 2025. p. 2025.10.08.25337572. doi:10.1101/2025.10.08.25337572

58. DeBoever C, Tanigawa Y, Lindholm ME, McInnes G, Lavertu A, Ingelsson E, et al. Medical relevance of protein-truncating variants across 337,205 individuals in the UK Biobank study. Nat Commun. 2018;9: 1612. doi:10.1038/s41467-018-03910-9

59. Trynka G, Hunt KA, Bockett NA, Romanos J, Mistry V, Szperl A, et al. Dense genotyping identifies and localizes multiple common and rare variant association signals in celiac disease. Nat Genet. 2011;43: 1193–1201. doi:10.1038/ng.998

60. Yates AD, Achuthan P, Akanni W, Allen J, Allen J, Alvarez-Jarreta J, et al. Ensembl 2020. Nucleic Acids Res. 2020;48: D682–D688. doi:10.1093/nar/gkz966

61. McLaren W, Gil L, Hunt SE, Riat HS, Ritchie GRS, Thormann A, et al. The Ensembl Variant Effect Predictor. Genome Biol. 2016;17: 122. doi:10.1186/s13059-016-0974-4

62. Karczewski KJ, Francioli LC, Tiao G, Cummings BB, Alföldi J, Wang Q, et al. The mutational constraint spectrum quantified from variation in 141,456 humans. Nature. 2020;581: 434–443. doi:10.1038/s41586-020-2308-7

63. Tanigawa Y, Qian J, Venkataraman G, Justesen JM, Li R, Tibshirani R, et al. Significant sparse polygenic risk scores across 813 traits in UK Biobank. PLoS Genet. 2022;18: e1010105. doi:10.1371/journal.pgen.1010105

64. Landrum MJ, Lee JM, Riley GR, Jang W, Rubinstein WS, Church DM, et al. ClinVar: public archive of relationships among sequence variation and human phenotype. Nucleic Acids Res. 2014;42: D980–5. doi:10.1093/nar/gkt1113

65. Venkataraman GR, Olivieri JE, DeBoever C, Tanigawa Y, Justesen JM, Dilthey A, et al. Pervasive additive and non-additive effects within the HLA region contribute to disease risk in the UK Biobank. bioRxiv. 2020. doi:10.1101/2020.05.28.119669

66. Chang CC, Chow CC, Tellier LC, Vattikuti S, Purcell SM, Lee JJ. Second-generation PLINK: rising to the challenge of larger and richer datasets. Gigascience. 2015;4: 7. doi:10.1186/s13742-015-0047-8

67. Inker LA, Eneanya ND, Coresh J, Tighiouart H, Wang D, Sang Y, et al. New creatinine- and cystatin C-based equations to estimate GFR without race. N Engl J Med. 2021;385: 1737–1749. doi:10.1056/NEJMoa2102953

68. Galinsky KJ, Bhatia G, Loh P-R, Georgiev S, Mukherjee S, Patterson NJ, et al. Fast Principal-Component Analysis Reveals Convergent Evolution of ADH1B in Europe and East Asia. Am J Hum Genet. 2016;98: 456–472. doi:10.1016/j.ajhg.2015.12.022

69. Mbatchou J, Barnard L, Backman J, Marcketta A, Kosmicki JA, Ziyatdinov A, et al. Computationally efficient whole-genome regression for quantitative and binary traits. Nat Genet. 2021;53: 1097–1103. doi:10.1038/s41588-021-00870-7

70. Bulik-Sullivan BK, Loh P-R, Finucane HK, Ripke S, Yang J, Schizophrenia Working Group of the Psychiatric Genomics Consortium, et al. LD Score regression distinguishes confounding from polygenicity in genome-wide association studies. Nat Genet. 2015;47: 291–295. doi:10.1038/ng.3211

71. Leonenko G, Baker E, Stevenson-Hoare J, Sierksma A, Fiers M, Williams J, et al. Identifying individuals with high risk of Alzheimer’s disease using polygenic risk scores. Nat Commun. 2021;12: 4506. doi:10.1038/s41467-021-24082-z

72. Bakulski KM, Vadari HS, Faul JD, Heeringa SG, Kardia SLR, Langa KM, et al. A non-APOE Polygenic score for Alzheimer’s disease and APOE-ε4 have independent associations with dementia in the Health and Retirement Study. medRxiv. 2020. doi:10.1101/2020.02.10.20021667

73. Skoog I, Kern S, Najar J, Guerreiro R, Bras J, Waern M, et al. A non-APOE polygenic risk score for Alzheimer’s disease is associated with cerebrospinal fluid neurofilament light in a representative sample of cognitively unimpaired 70-year Olds. J Gerontol A Biol Sci Med Sci. 2021;76: 983–990. doi:10.1093/gerona/glab030

74. Ware EB, Faul JD, Mitchell CM, Bakulski KM. Considering the APOE locus in Alzheimer’s disease polygenic scores in the Health and Retirement Study: a longitudinal panel study. BMC Med Genomics. 2020;13: 164. doi:10.1186/s12920-020-00815-9

75. Purcell S, Neale B, Todd-Brown K, Thomas L, Ferreira MAR, Bender D, et al. PLINK: a tool set for whole-genome association and population-based linkage analyses. Am J Hum Genet. 2007;81: 559–575. doi:10.1086/519795

