## Supplementary Figures 1-7 for "PRISM: ancestry-aware integration of tissue-specific genomic annotations enhances the transferability of polygenic scores"

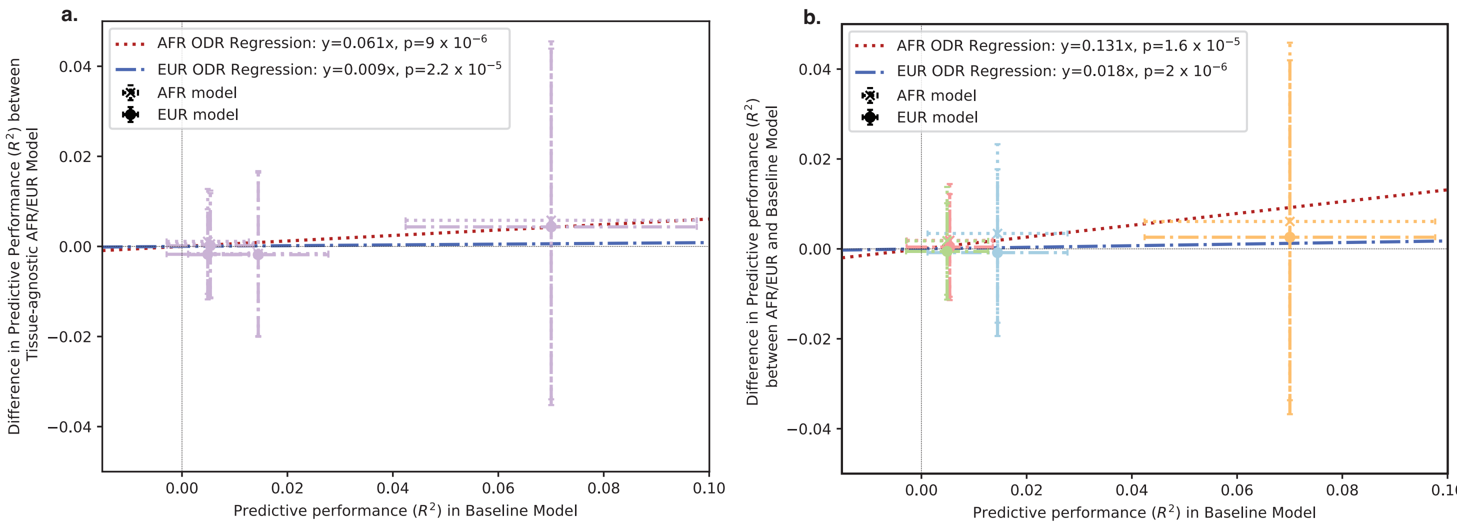


**Supplementary Fig. 1. Average improvement in prediction accuracy across four select traits**

a. We compare prediction accuracy of ancestry-matched (AFR, red regression line), and ancestry-mismatched (EUR, blue regression line) PRISM model against that of the baseline model using tissue non-specific annotations. The x-axis represents the predictive performance of the baseline model, evaluated in $R^2$, while the y-axis shows the difference in predictive performance between the PRISM models and the baseline model, also evaluated in $R^2$. The average improvement is quantified by applying orthogonal distance regression, with the improvement represented by the fitted coefficient (Methods). We observe a greater fitted coefficient, $\beta_{AFR}$=0.131 for ancestry-matched PRISM compared to $\beta_{EUR}$=0.018 for ancestry-mismatched PRISM.

b. We compare the predictive accuracy of ancestry-matched (AFR, red regression line), and ancestry-mismatched (EUR, blue regression line) PRISM model against that of the baseline model using tissue-matched annotations. The x-axis represents the predictive performance of the baseline model, evaluated in $R^2$, while the y-axis shows the difference in predictive performance between the PRISM models and the baseline model, also evaluated in $R^2$. The average improvement is quantified by applying orthogonal distance regression, with the improvement represented by the fitted coefficient (Methods). Scatter points are color-coded by tissue-specificity. We observe a greater fitted coefficient, $\beta_{AFR}$=0.061 for ancestry-matched PRISM compared to $\beta_{EUR}$ = 0.009 for ancestry-mismatched PRISM.

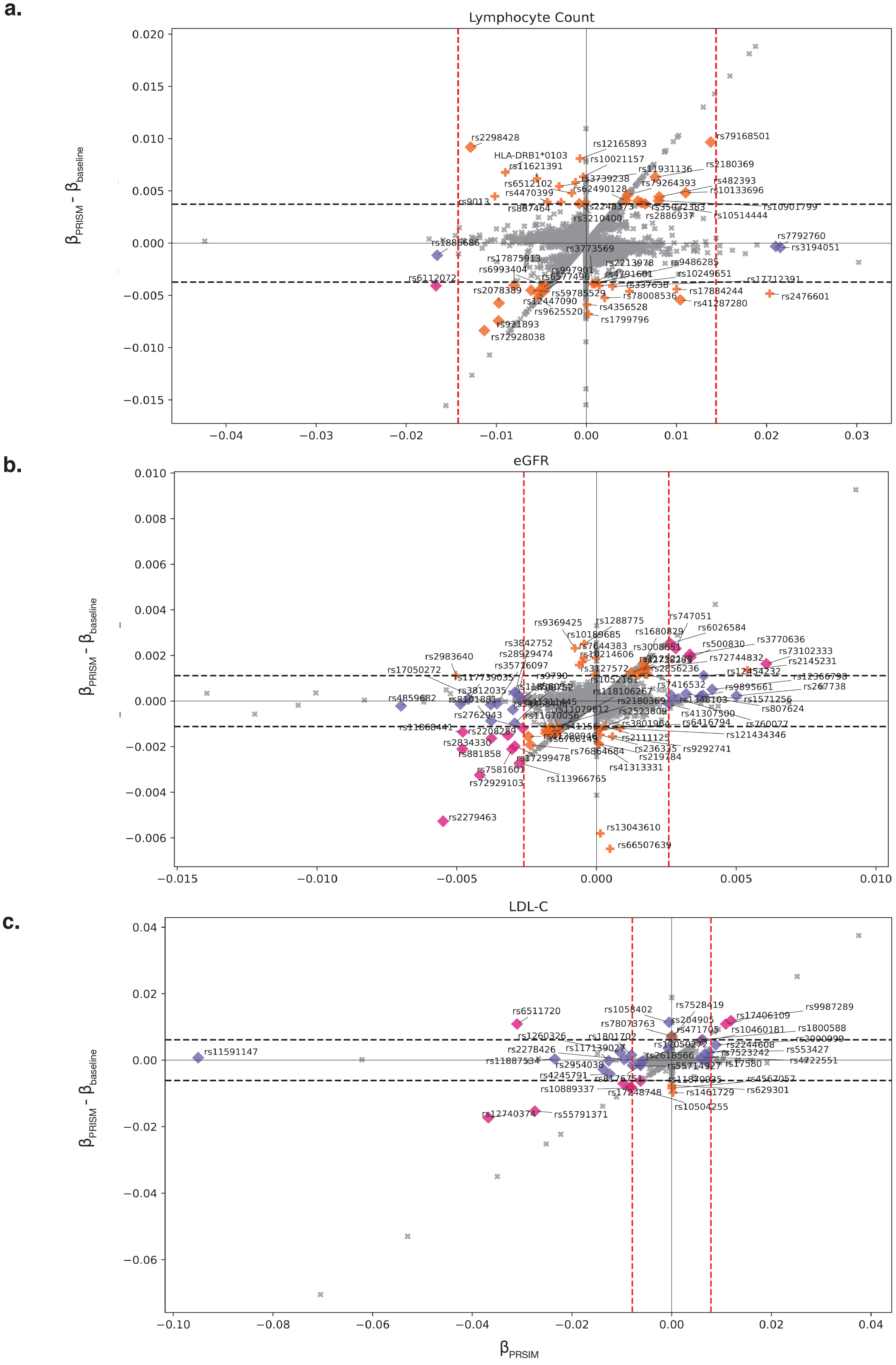


**Supplementary Fig. 2. PRISM-identified biologically relevant variants and their functional consequences in the regulatory landscape.**

We group variants into four categories: A, B, C, and D, with decreasing priority (colors). The grouping of variants is determined using three criteria: i. the absolute value of the variant effect size (red dashed vertical line), ii. the absolute value of the difference in effect sizes between PRISM and the baseline model (black dashed horizontal line), and iii. whether the variants are prioritized according to their continuous cV2F scores (diamond: prioritized, cross: not prioritized)

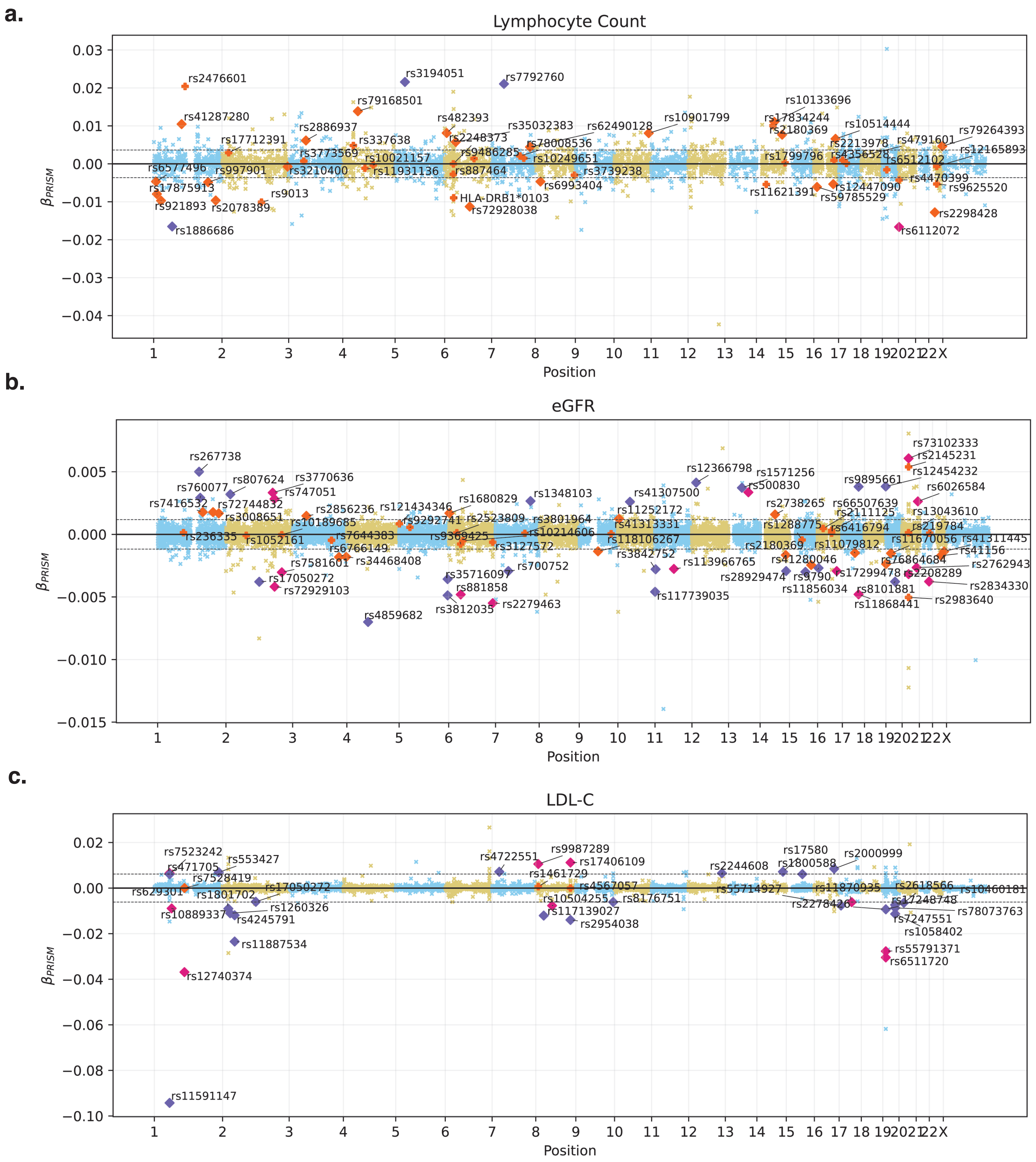


**Supplementary Fig. 3. Genetic variants' effect derived from ancestry- and tissue-matched PRISM across the genome of select traits.**
We show PGS effect sizes of genetic variants from the ancestry- and tissue- matched PRISM models for lymphocyte count, eGFR, and LDL-C. Variants are colored by group attribute.

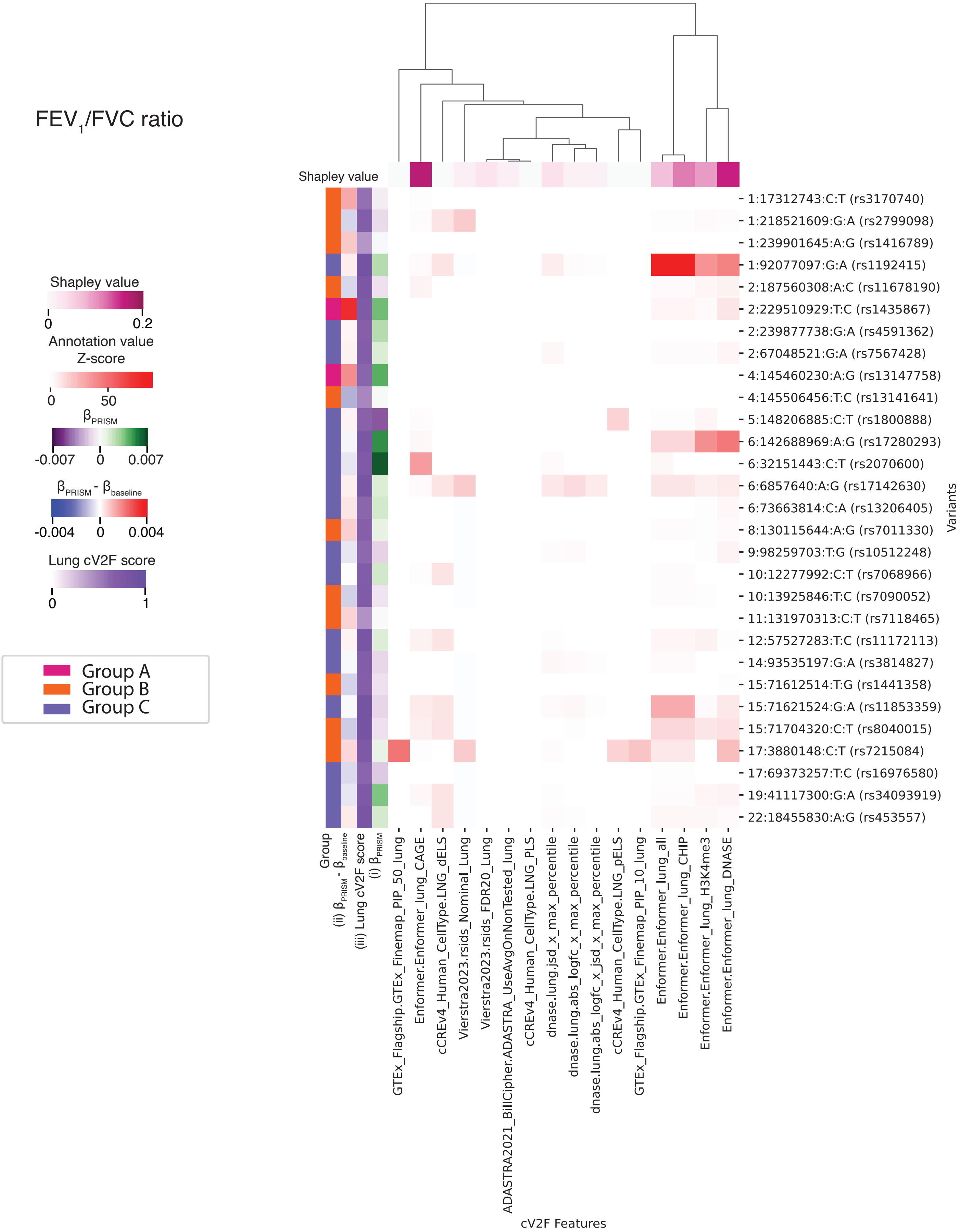


**Supplementary Fig. 4. Contribution of lung annotations to improved PGS transferability for FEV1/FVC ratio (full Z-score scale)**

We show the Z-scores of annotation values (x-axis) for grouped variants across the genome (y-axis). We show the SHAP value for each annotation at the top of the heatmap. On the left of the heatmap, we display the group attribute for each variant, the PGS effect size difference between the PRISM and baseline model, and the continuous lung cV2F score in order. JSD stands for Jensen–Shannon Divergence.

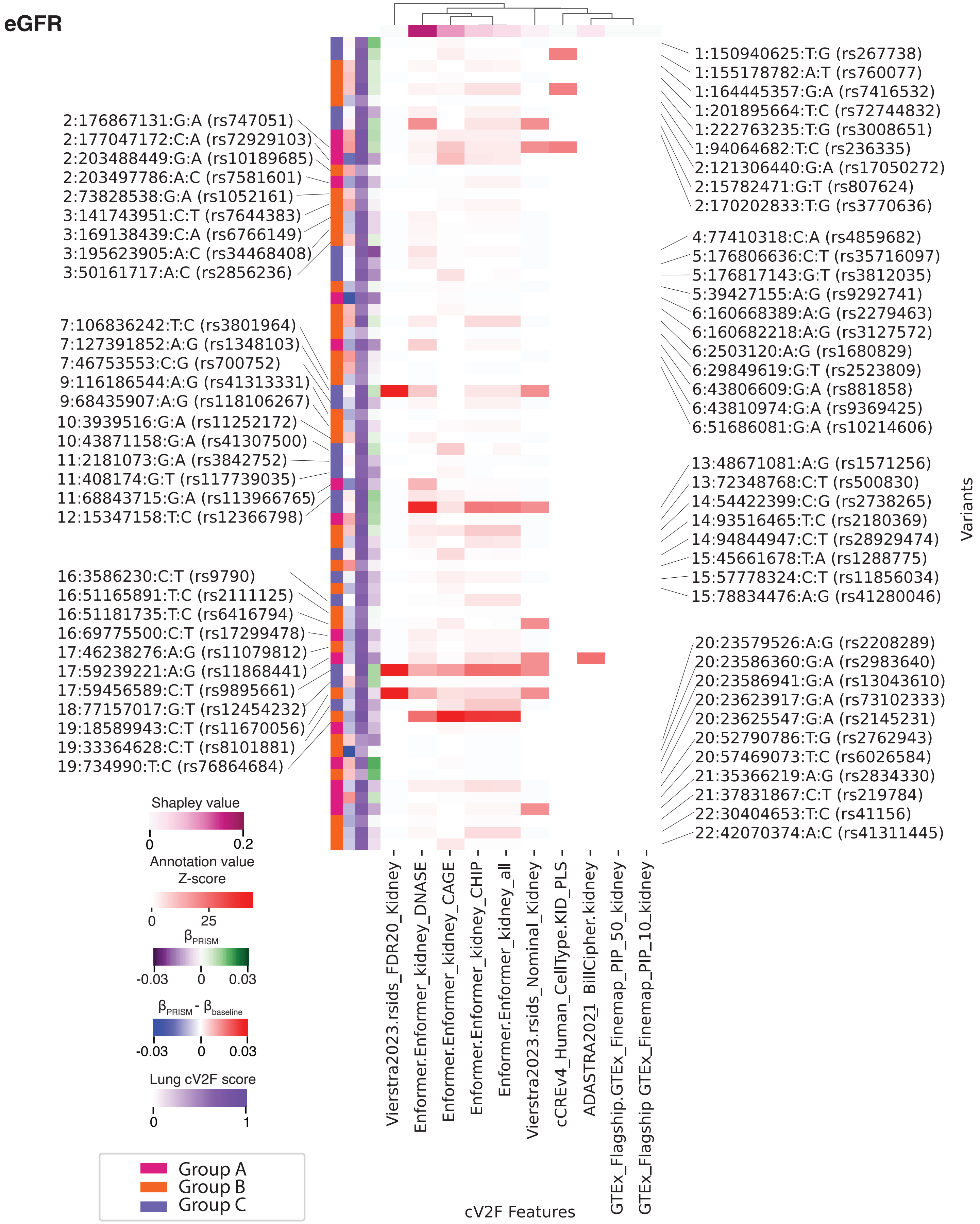


**Supplementary Fig. 5. Contribution of kidney annotations to improved PGS transferability for eGFR (full Z-score scale)**

We show the Z-scores of annotation values (x-axis) for grouped variants across the genome (y-axis). We show SHAP value for each annotation at the top of the heatmap. On the left of the heatmap, we display the group attribute for each variant and criterion ii, iii, and i in order. JSD stands for Jensen–Shannon Divergence.

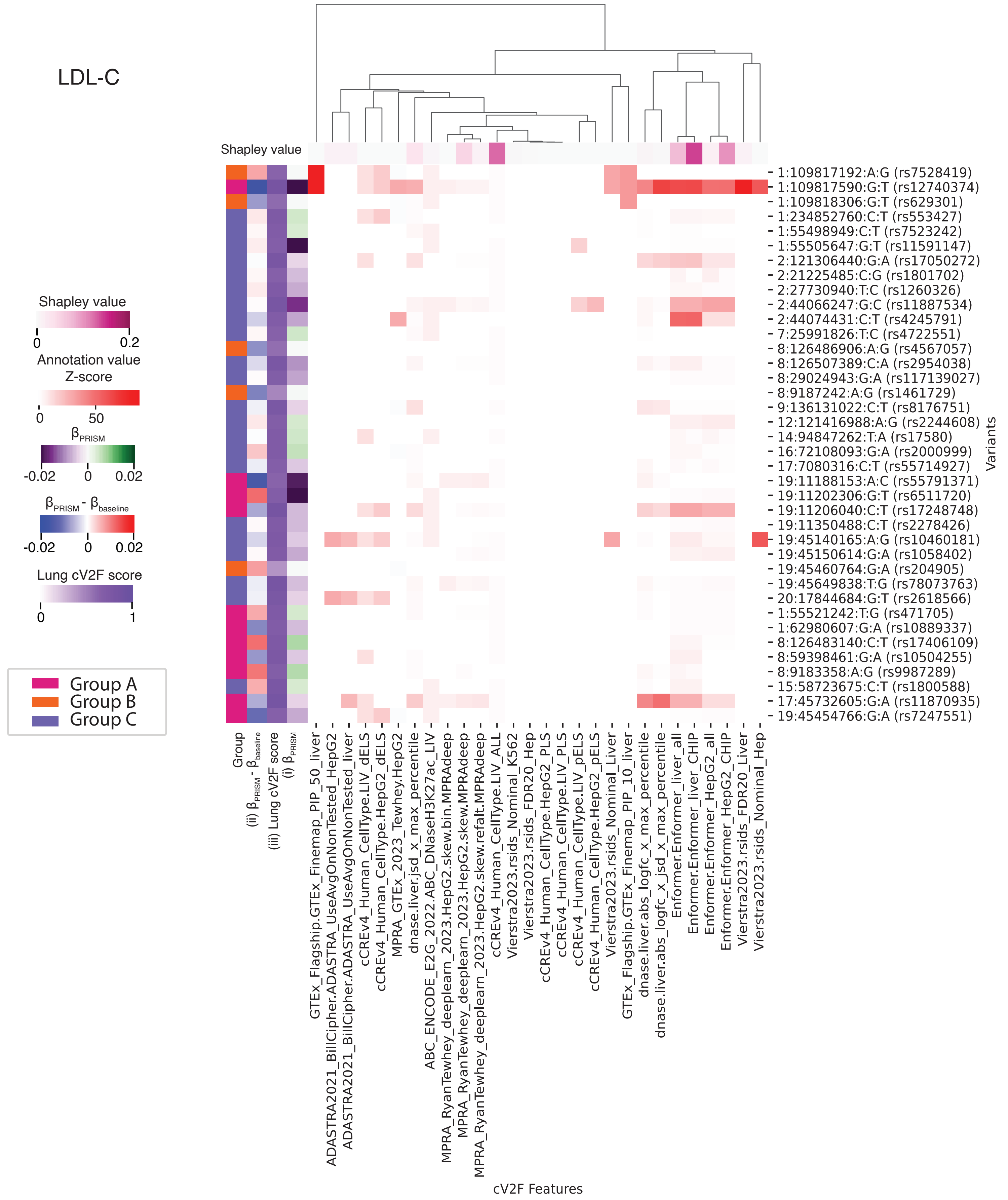


**Supplementary Fig. 6. Contribution of liver annotations to improved PGS transferability for LDL-C (full Z-score scale)**

We show the Z-scores of annotation values (x-axis) for grouped variants across the genome (y-axis). We show the SHAP value for each annotation at the top of the heatmap. On the left of the heatmap, we display the group attribute for each variant and criteria ii, iii, and i in order. JSD stands for Jensen–Shannon Divergence.

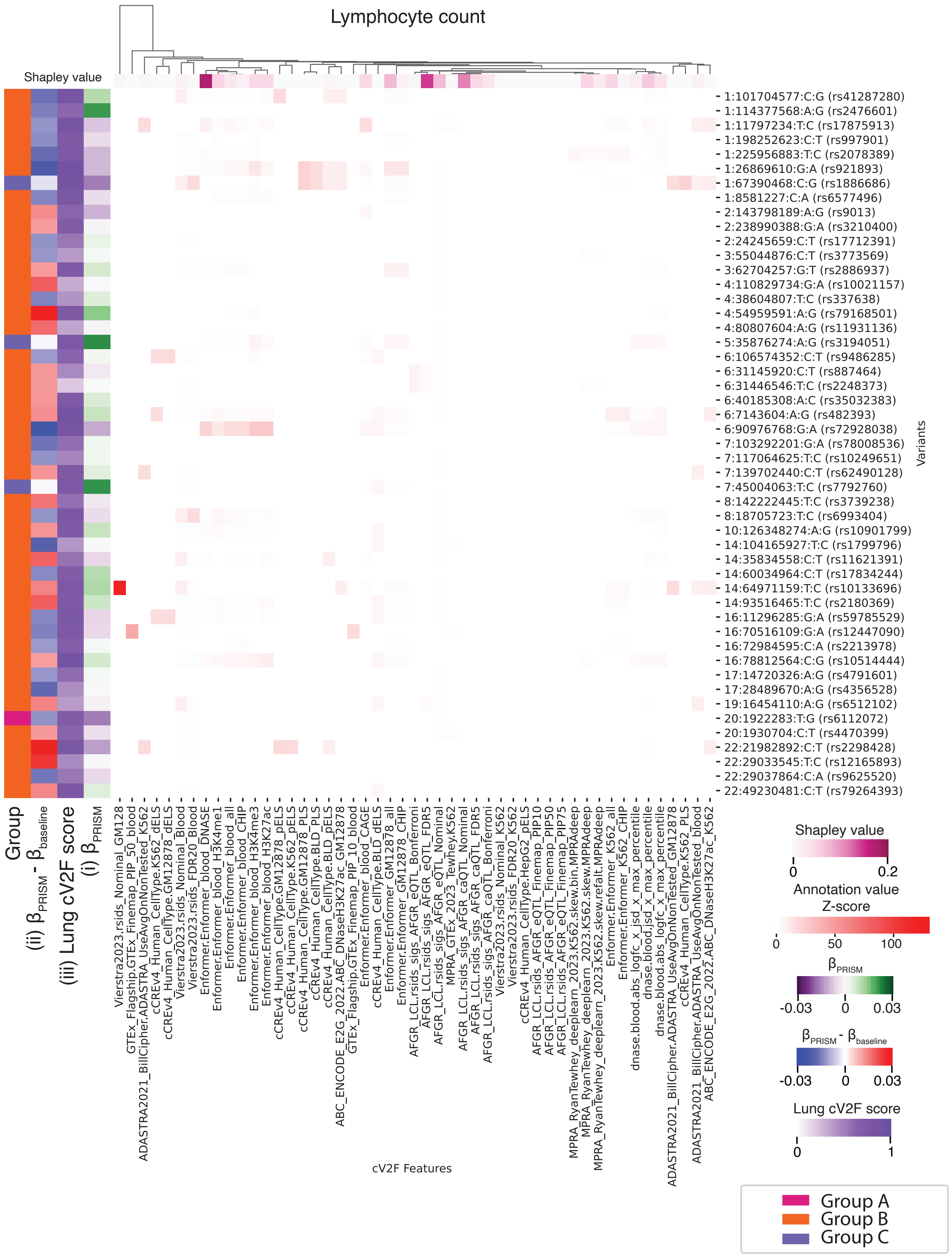


**Supplementary Fig. 7. Contribution of blood annotations to improved PGS transferability for lymphocyte count (full Z-score scale)**

We show the Z-scores of annotation values (x-axis) for grouped variants across the genome (y-axis). We show the SHAP value for each annotation at the top of the heatmap. On the left of the heatmap, we display the group attribute for each variant and criteria ii, iii, and i in order. JSD stands for Jensen–Shannon Divergence.
